# A pervasive large conjugative plasmid mediates multispecies biofilm formation in the intestinal microbiota increasing resilience to perturbations

**DOI:** 10.1101/2024.04.29.590671

**Authors:** Leonor García-Bayona, Nathania Said, Michael J. Coyne, Katia Flores, Nazik M. Elmekki, Madeline L. Sheahan, Adriana Gutierrez Camacho, Kathryn Hutt, Fitnat H. Yildiz, Ákos T. Kovács, Matthew K. Waldor, Laurie E. Comstock

## Abstract

Although horizontal gene transfer is pervasive in the intestinal microbiota, we understand only superficially the roles of most exchanged genes and how the mobile repertoire affects community dynamics. Similarly, little is known about the mechanisms underlying the ability of a community to recover after a perturbation. Here, we identified and functionally characterized a large conjugative plasmid that is one of the most frequently transferred elements among Bacteroidales species and is ubiquitous in diverse human populations. This plasmid encodes both an extracellular polysaccharide and fimbriae, which promote the formation of multispecies biofilms in the mammalian gut. We use a hybridization-based approach to visualize biofilms in clarified whole colon tissue with unprecedented 3D spatial resolution. These biofilms increase bacterial survival to common stressors encountered in the gut, increasing strain resiliency, and providing a rationale for the plasmid’s recent spread and high worldwide prevalence.

## INTRODUCTION

Bacteroidales is the dominant order of Gram-negative bacteria of the human gut microbiota^1^, with species of the genera *Bacteroides*, *Phocaeicola* and *Parabacteroides* dominating in individuals living in industrialized populations^2^. These bacteria are remarkably stable, with many strains colonizing for decades, although some are occasionally lost or replaced^3,4^. Strains that stably colonize the human gut must be resilient to many varying environmental factors of this ecosystem including nutrient fluxes due to diet, microbial compositional fluxes, antagonism, bouts of inflammation, xenobiotics, and particularly in the industrialized world, the consumption of antibiotics. The mechanisms allowing for persistence despite varying environmental conditions are still not well understood. More broadly, the gut microbiome displays substantial resilience to moderate perturbations, whereas harsher changes can lead to permanent compositional shifts^5–7^. Addressing these questions becomes more pressing as modern lifestyles and antibiotic overuse has led to decreased species diversity and increasingly abundant microbiome-associated pathologies^8^.

Horizontal gene transfer (HGT) is pervasive within the human gut microbiota, especially in industrialized populations, such that each person’s microbiota becomes “personalized”^9,10^. Recent efforts have systematically catalogued the mobilome in human gut microbiomes, yet the physiological roles of a large proportion of cargo genes carried on these mobile genetic elements (MGE) remains unknown or only broadly classified^10–13^. HGT events are prevalent between species within a phylum and rare between species of different phyla^10,11,14^. While cargo genes in many MGEs are globally distributed, the genetic architectures of MGEs are not generally conserved^10,15^. Frequent recombination and gene loss/gain events lead to shuffling of cargo and mobilization genes^10,11,15,16^. Industrialized and non-industrialized populations differ significantly in the composition of their gut microbiota, with limited gene flow between them^10,15,17^.

We previously analyzed MGEs that had transferred between co-resident Bacteroidales species (from the same individual) in four cohorts from the USA^16,18,19^, UK^20^ and China^21^. We identified 74 MGE that spread through multiple species within individual gut microbiomes^16^. The vast majority of these MGE were present in only one or a few subjects and most showed evidence of frequent recombination so that they did not have a conserved architecture between individuals. Three notable exceptions were a ubiquitous, well-studied conjugative transposon harboring a tetracycline resistance gene,^22–24^ and two MGEs carrying Type VI secretion system loci^25^ involved in bacterial antagonism. We also identified a fourth such element, a 98 kb plasmid with a conserved architecture that we designated as pMMCAT, which also spread among multiple Bacteroidales species in many communities^16^.

Our studies here show that pMMCAT encodes genes that mediate the formation of multispecies biofilms in the colon. Biofilms are aggregates of microbial cells embedded within a protective extracellular matrix composed of various molecules, including polysaccharides, proteins, and DNA^26^. Biofilms play an important role in survival from environmental insults^26,27^ and enable beneficial interactions such as syntrophy, DNA exchange, and break-down of complex nutrient particles.^26,27^. Gut mucosal biofilms have been studied in pathogens and in colorectal cancer^28–30^, but biofilms formed by the microbiota in the healthy gut are less studied^30^. Two gut Bacteroidales strains were shown to form mixed-species biofilms *in vitro* and to engage in metabolite sharing^31^. We demonstrate that pMMCAT-mediated biofilm formation increases bacterial survival under common environmental stressors, such as antibiotics, leading to improved community resilience to perturbations in the gut.

## RESULTS

### The genetic repertoire of pMMCAT

Our initial report of the discovery of the pMMCAT large conjugative plasmid showed that it is exceptional due its spread to many (up to five) different Bacteroidales species in the gut microbiomes of five different human subjects analyzed, with a highly conserved architecture throughout its 98 kb sequence^16^. Analysis of read counts in our whole-genome sequencing of 16 strains carrying pMMCAT indicates that in liquid culture this plasmid is maintained at one to three copies per cell and our long-read sequencing indicates that it is a circular molecule (Fig. S1A-C). pMMCAT carries a full complement of genes encoding type IV conjugation machinery and is therefore not dependent on the machinery of other mobile elements for its transfer (Fig. 1A). It has two notable loci that suggested it may mediate biofilm formation, therefore we named this plasmid pMMCAT (<u>p</u>lasmid-<u>M</u>ediated <u>M</u>icrobiome <u>C</u>ommunity <u>At</u>tachment). The first locus is 24-kb and contains genes for the synthesis of an extracellular polysaccharide (EPS) with nine glycosyltransferase genes, suggesting a large glycan repeat unit. EPS is a crucial component of the extracellular matrix of biofilms^32^. The synthesis of this EPS is phase variable, dictated by two adjacent invertible promoters with a tyrosine DNA recombinase family invertase encoded upstream (Fig. 1A). pMMCAT also harbors an Mfa-like short adhesive fimbriae locus, with an adjacent LuxR-like transcriptional regulator, which often respond to quorum sensing auto-inducers or other small molecules^33^. In biofilms, fimbriae are important for cell surface attachment as well as cell-cell adhesion^34^. Other genes carried by pMMCAT include a putative anti-CRISPR system, as well as five toxin-antitoxin “addiction module” pairs that potentially promote plasmid maintenance. A 4.8-kb region encodes two radical S-adenosyl-L-methionine enzymes and a metallo-β-lactamase-fold protein. These products may confer resistance to toxic molecules, mediate post-translational protein modifications, or be involved in the synthesis of biologically active small molecules^35,36^.

**Figure 1.**
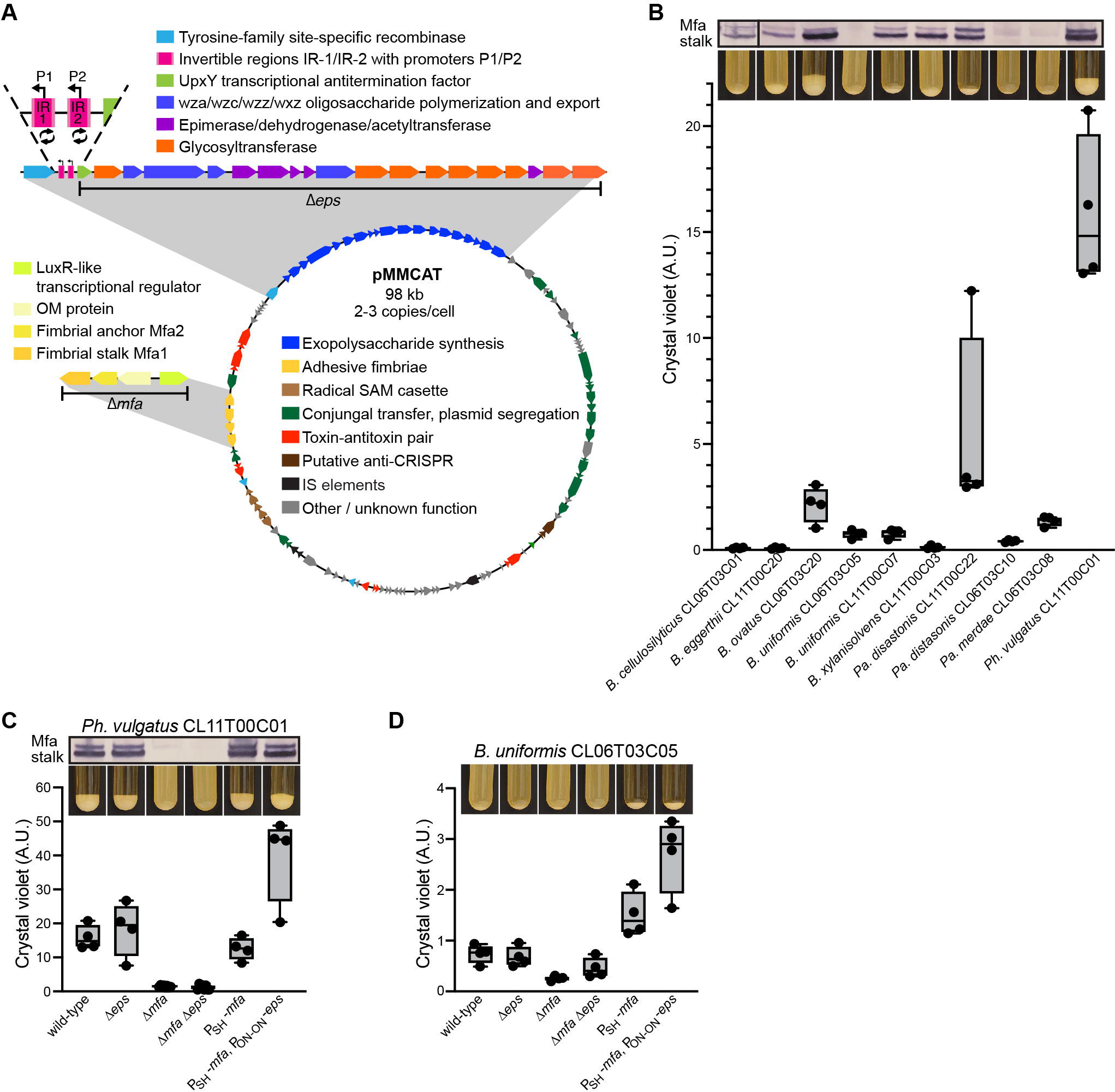
pMMCAT mediates biofilm formation. **A.** Schematic of pMMCAT loci. IR: Invertible region. P1/P2: promoter regions **B.** Top panel: Immunoblot of cells of the specified strain with antibody against the Mfa fimbriae stalk protein (INE91_04914 in PhvCL11). Middle panel: Bacterial aggregation of the specified strain grown in stationary culture. Bottom panel: Quantification of crystal violet staining for biofilm formation in 24-well plates (four biological replicates shown as individual data points, whisker boxes represent quartiles). **C.** Quantification of biofilm formation by PhvCL11 mutants (regions deleted in *mfa* and *eps* mutants shown in panel A). Top, middle, and bottom panels as in panel B. **D.** Quantification of biofilm formation by *B. uniformis* CL06T03C05 mutants. Top and bottom panels as in panel B.

### pMMCAT mediates biofilm formation

Due to the presence of the fimbriae and EPS loci, we predicted that pMMCAT may confer the ability to form biofilms. Most *Bacteroides*, *Phocaeicola* and *Parabacteroides* strains we have cultured do not spontaneously aggregate when grown under static conditions in liquid culture^37,38^. We evaluated the ability of ten wild-type strains of eight different species harboring pMMCAT to aggregate and form biofilms. Upon entry into early stationary phase, four of these strains formed strong cell-cell aggregates that settled to the bottom of the tube (Fig. 1B), with very few planktonic cells remaining. The other six strains harboring pMMCAT also formed visible cell-aggregates at the bottom of the tube, but with variable amounts of planktonic cells. Although very tightly associated, these cell aggregates lacked any observable attachment to the glass culture tube as is observed in many biofilm-forming bacteria. There was a strong correlation between the degree of aggregation in culture and the level of Mfa produced by the bacteria (Fig 1B, Sig S1D). However, *B. cellulosilyticus* CL06T03C01 produced a slight amount of Mfa but did not exhibit any aggregation, and Mfa is highly produced in *B. xylanisolvens* CL11T00C03 yet a large fraction of cells remained planktonic, indicating that additional factors influence the phenotype. We used crystal violet staining to quantify biofilm formation in 24-well plates, by carefully removing the planktonic fraction and staining the aggregated cells at the bottom of the well. As in the glass culture tubes, the bacteria did not attach to tissue-culture treated polystyrene but formed a robust pellet at the bottom of the well. Seven of the ten strains displayed some amount of pellet formation, with *Phocaeicola vulgatus* CL11T00C01 (PhvCL11) forming the largest aggregates (Fig. 1B). This variation across strains is not surprising given that pMMCAT is present in very different genomic contexts in these strains, where regulatory networks may vary significantly.

To conclusively determine that the *mfa* and *eps* loci encode fimbriae and EPS, respectively, and that they are involved in the biofilm phenotype, we generated *eps* and *mfa* deletion mutants in the strongest biofilm former isolate, PhvCL11, as well as in a strain that displayed limited biofilm formation (*Bacteroides uniformis* CL06T03C05, BuCL06). In PhvCL11, a deletion of *mfa* completely abrogated the aggregative phenotype, as did a double *ΔepsΔmfa* deletion (Fig.1A). In contrast, a deletion of the complete *eps* locus had no impact on the amount of aggregation (Fig. 1C, Fig. S1E). However, even if the aggregative property remains, the protective matrix feature of the biofilm is likely reduced in the *Δeps*, leading to an altered biofilm.

Interestingly, placing the *mfa* operon under a strong constitutive promoter in PhvCL11 did not change the amount of pellet formed in the biofilm assay, whereas it was increased when coupled with constitutive expression of the *eps* locus by locking its invertible promoters in the “on” orientation. This result suggests that in static cultures, not all cells are expressing the *eps* locus. This type of population bet-hedging in the Bacteroidales has been reported for phase-variable loci encoding capsular polysaccharides and other surface molecules^39–42^. To evaluate if BuCL06 has the potential to form robust biofilms, we placed *mfa* behind a strong constitutive promoter and locked the *eps* promoter regions in the “on” orientation (Fig. 1D). Expression of *mfa* was sufficient to cause very strong aggregation. When coupled with constitutively expressed *eps*, biofilm formation was significantly increased. These results indicate that both loci contribute to pMMCAT-mediated biofilm formation and are differentially expressed in different strains grown in liquid culture. To determine if the secreted polysaccharide remains primarily envelope-associated like a capsular polysaccharide or forms a more diffuse matrix in the biofilm (an EPS), we performed an immunoblot of culture supernatants using polyclonal antibodies against the EPS. It showed that this secreted polysaccharide is detectable and is of high molecular weight (Fig. S1F).

We used laser scanning confocal microscopy (LSCM) to visualize the biofilms formed by PhvCL11. When PhvCL11 wild type strains producing either GFP or mCherry were co-cultured at equal ratios, the pellet biofilm was composed of approximately equal amounts of both (Fig 2A, B). In contrast, the PhvCL11 *ΔepsΔmfa* mutant did not produce a biofilm. Consistent with the crystal violet quantification, a deletion of the *mfa* locus was sufficient to remove the ability to form a biofilm, whereas a *Δeps* mutant still formed aggregates. Interestingly, the PhvCL11 *ΔepsΔmfa* mutant was able to incorporate into a PhvCL11 wild-type biofilm, indicating that, while necessary for establishing a biofilm when alone, these two loci are not required to join an established biofilm. Similarly, a *Δmfa* mutant was able to incorporate into a wild-type biofilm, although it was significantly depleted in these biofilms. Likewise, co-culture of *Δeps* with *ΔepsΔmfa* resulted in an enrichment of the *Δeps* in the cell aggregates.

**Figure 2.**
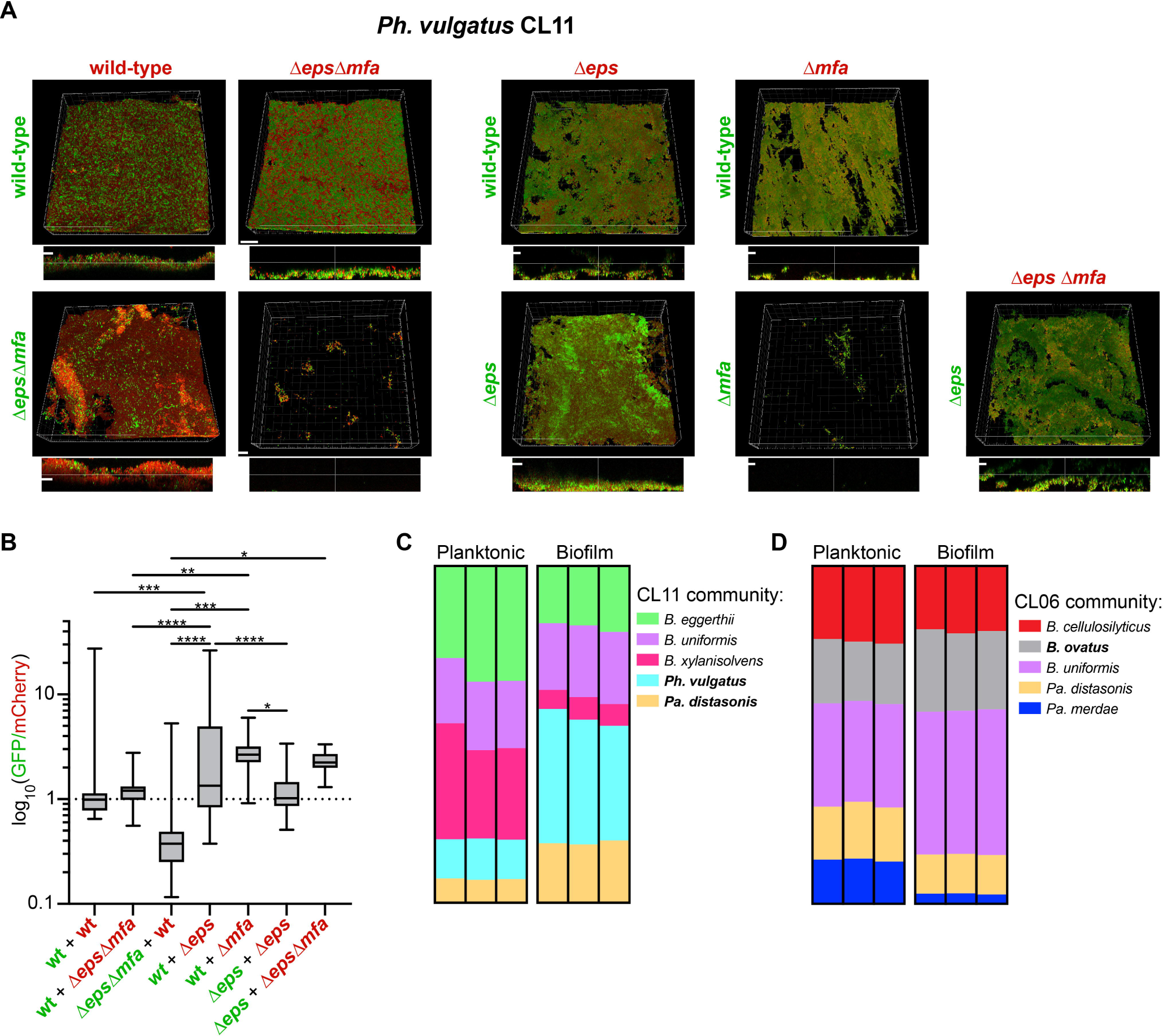
pMMCAT-mediated biofilms in liquid culture. **A.** Laser scanning confocal microscopy of biofilms formed by the specified strains, expressing either GFP (green) or mCherry (red). Scale bar: 100 µm. Top: Blend 3D projection, Bottom: cross-section along the middle vertical and horizontal planes. **B.** Quantification of the abundance of each strain in the specified pairwise co-cultures, as indicated in panel A. For each combination, a minimum of 16 frames from a minimum of 3 independent biological replicates were analyzed. **C-D.** Composition of the biofilm and planktonic fractions of a liquid co-culture of the specified strains (3 independent biological replicates). Abundances were determined by shotgun metagenomic sequencing (3 biological replicates are shown side to side).

Co-cultures under static conditions of five pMMCAT-harboring species isolated from the same person (CL11) formed a robust pellet biofilm that did not dissociate even with vigorous shaking, while a fraction of cells remained planktonic (Video S1). We used high-throughput sequencing to quantify the relative abundance of each species in the pellet biofilm and in the planktonic fraction (Fig. 2C). Although all five species could be detected in both the biofilm and planktonic fractions, many were enriched or depleted in one or the other. The two strains that form biofilms in pure culture (Fig. 1B), PhvCL11 and *P. distasonis*, were enriched in the biofilm whereas *B. xylanisolvens*, which failed to form a biofilm in pure culture, was almost completely excluded from the mixed species biofilm and was the most abundant planktonic species. *B. uniformis* and *B. eggerthii*, which form biofilms in monoculture, were detected in the community biofilm and in the planktonic fraction. These results support the conclusion that pMMCAT mediates *bona fide* multispecies biofilms. Interestingly, in a different five-member pMMCAT containing consortium (CL06), where only one strain forms biofilms in monoculture (*B. ovatus*), the biofilm fraction became dominated by *B. uniformis*, a species that failed to produce biofilm in monoculture (Fig. 2D), suggesting cross-species regulation.

To study the spatial organization of this biofilm in the colon, we used MiPACT (Microbial identification after Passive CLARITY Technique) followed by Hybridization Chain Reaction (HCR)^43,44^. In short, gnotobiotic mice were colonized with PhvCL11 for two weeks after which the entire colon was extracted, fixed, and embedded in a hydrogel. Through a series of steps, the colon become optically transparent (Fig S2A), and probes were hybridized to a highly expressed mRNA (Fig S2B) and imaged in z-stacks using LSCM. We used probes against *mCherry* to visualize PhvCL11 cells, and probes to detect the *eps* locus mRNA to evaluate where cells are producing this key component of the biofilm matrix. There were no major differences in overall bacterial spatial distribution in mice monocolonized with wild-type PhvCL11 or with the Δ*eps*Δ*mfa* isogenic mutant (Fig. 3, Videos S2-S4). Notably, no cell aggregates were observed directly attached to the mucosa, as occurs with *Fusobacterium nucleatum* implicated in colorectal cancer^45,46^. Interestingly, we observed little evidence of *eps* expression above background levels in mice except potentially in small clusters associated with plant material (Fig. 3A, Fig. S2C, Video S2). Next, we treated mice with metronidazole to analyze *eps* expression during bacterial stress. This stressor was selected because, as discussed later, the *eps* and *mfa* loci are strongly induced when mice are treated with metronidazole, increasing bacterial survival during this treatment. Our imaging analyses showed that during metronidazole treatment, the *eps* locus is highly expressed in the bacteria at the mucus layer, and on the surface and internal compartments of some food particles (Fig. 3B, Fig. S2D, Video S3). In regions of the colon with no fecal pellet, few bacteria were detected, mostly close to the mucus layer, with no *eps* expression above background Fig. 3B, Fig. S2D, Video S3). Only a faint signal was detected in mice monocolonized with the *ΔepsΔmfa* mutant, in the presence or absence of metronidazole (Fig. 3C, Video S4), confirming the high specificity of the *eps* probes. Therefore, this exopolysaccharide key component of colonic biofilms is induced in mice by metronidazole treatment and *eps* expressing bacteria are primarily localized to the surface of the fecal pellet at the mucus layer and inside some food particles.

**Figure 3.**
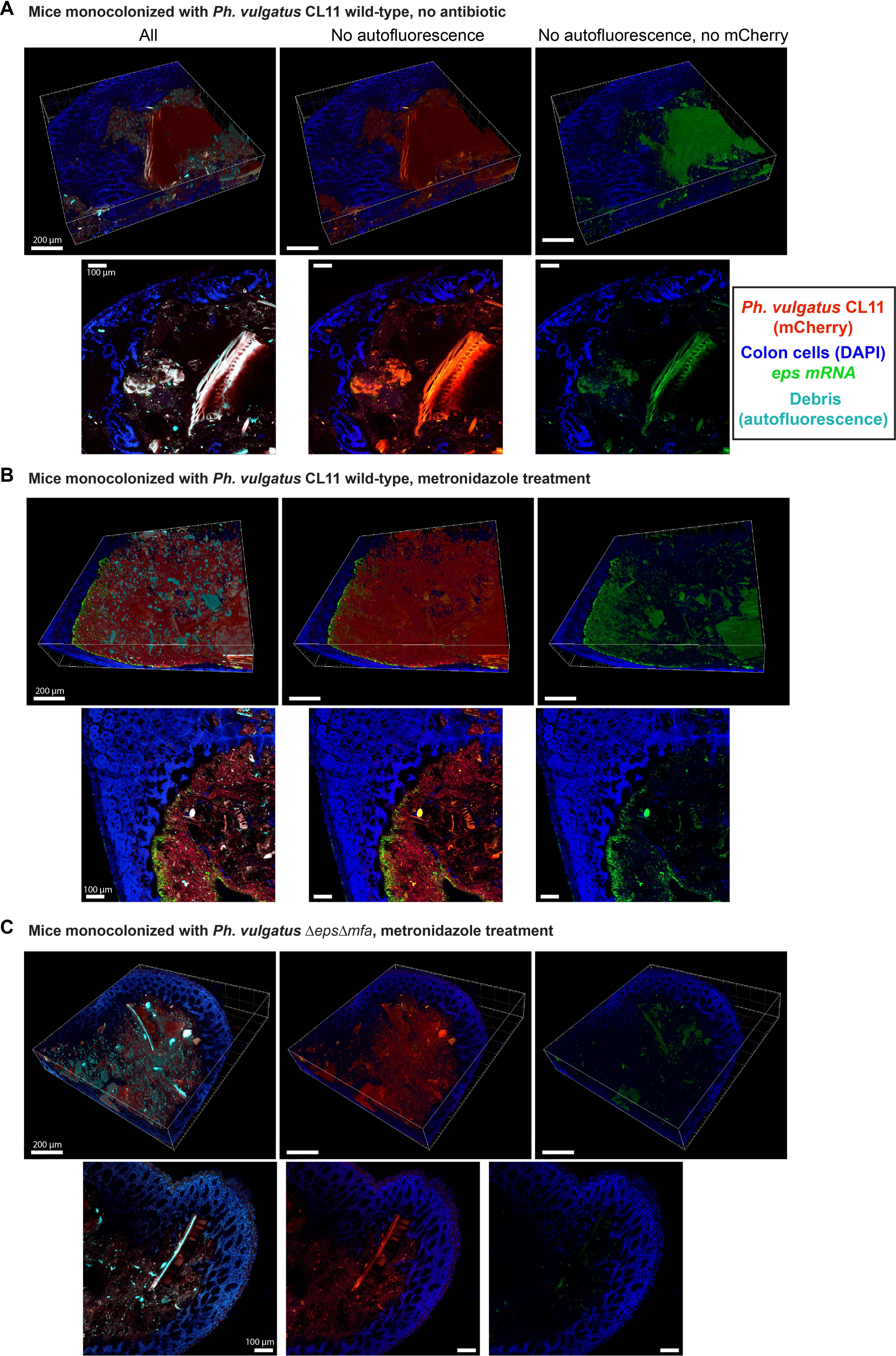
MiPACT-HCR 3D imaging of pMMCAT biofilms in the colon. Mice were monocolonized with PhvCL11 (constitutively expressing *mCherry*) and colons were imaged prior to **(A)** and during metronidazole treatment **(B)**. Mice monocolonized with the *ΔepsΔmfa* biofilm null mutant were used as control **(C)**. For each set of images, the top row shows the 3D Blend Projection in Imaris (scale bar 200 µm), and the bottom row shows a representative slice along the z-stack (scale bar 100 µm). DAPI (shown in blue) was used to stain host cell nuclei and bacterial cells, while autofluorescence of fecal pellet debris was imaged in the far red (shown in cyan). HCR probes against *mCherry* (shown in red) were used to detect all transcriptionally active PhvCL11 cells, while probes against the *eps* locus (shown in green) were used as a proxy for expression of biofilm genes. Representative images of one mouse are shown, but at least three mice per strain per condition were evaluated.

### pMMCAT-mediated biofilms increase resilience to stressors

Biofilms are often involved in protection against environmental insults. We evaluated the protective capabilities of pMMCAT-mediated biofilms to a panel of antibacterial stressors that these bacteria encounter in the intestine, including the clinically relevant antibiotics enrofloxacin, metronidazole and meropenem, the bile salt deoxycholate, human antimicrobial peptide LL-37 and the bacteriocin Bacteroidetocin A^47^. Bacterial survival was compared after stressors were added to an established static culture biofilm of wild-type PhvCL11 or to the *ΔepsΔmfa* mutant grown for the same amount of time (Fig. 4A). Biofilms protected against metronidazole at concentrations below 5 µg/ml and against low concentrations of enrofloxacin but not against meropenem. pMMCAT-mediated biofilms also increased survival to deoxycholate and to LL-37. Finally, the biofilm null mutant was more sensitive to Bacteroidetocin A at 4 ng/ml. Altogether, these results indicate that pMMCAT biofilms confer substantial protection to a diverse array of antimicrobial agents with different mechanisms of action *in vitro*.

**Figure 4.**
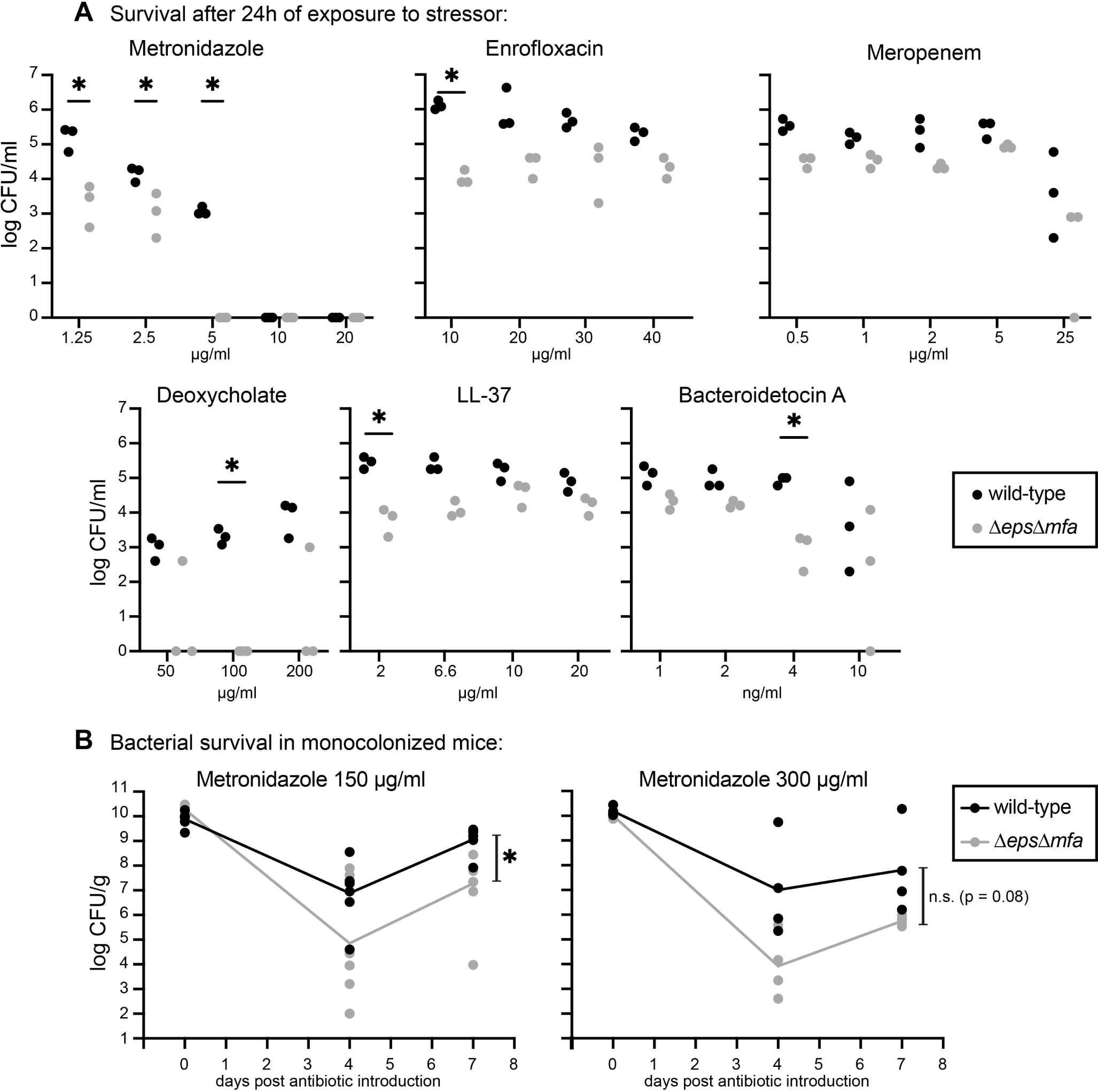
pMMCAT-mediated biofilms increase tolerance to chemical insults. **A.** Survival of PhvCL11 (wild-type of *ΔepsΔmfa*) to the listed stressors after 24 hours (three independent replicates). * indicates p < 0.05 (corrected for multiple unpaired t-tests using the Holm-Šídák method). **B**. Gnotobiotic mice were monocolonized for two weeks with either wild-type or *ΔepsΔmfa* PhvCL11 and treated with metronidazole at the specified concentration in the drinking water. For 150 µg/ml, six mice per treatment were used, housed in groups of 3. For 300 µg/ml, four mice per treatment were used, housed in pairs. Survival of the bacteria in the fecal pellet was evaluated by plating. Line connects geometric means of each treatment across timepoints. * indicates p < 0.05 comparison between strains in a two-way ANOVA. n.s. = non-significant.

To determine if pMMCAT confers similarly enhanced protection in the colon, we used a mouse model to quantify bacterial survival to metronidazole treatment (Fig. 4B). Mice were monocolonized with either wild-type PhvCL11 or the *ΔepsΔmfa* mutant, and metronidazole was added to the drinking water after two weeks of colonization. The metronidazole concentrations correspond to 0.15 and 0.30 times the concentration generally used to completely deplete Bacteroidales species^48^. Prior to antibiotic treatment, both strains colonized at comparable levels (Fig 4B). At both metronidazole concentrations, there were rapid decreases in the PhvCL11 CFU in the feces by day 4, followed by recovery. At 150 µg/ml, the initial drop in CFU was significantly greater in the *ΔepsΔmfa* mutant than in wild type. This trend was also observed at 300 µg/ml, but there was large mouse-to-mouse variation. In the context of a complex community, such variation in the magnitude of a decline can determine whether a strain is lost or is later able to recover. Therefore, the pMMCAT-mediated increase in tolerance to metronidazole and other stressors may have important impacts on strain resilience in a complex community.

We further explored the ability of pMMCAT to provide resilience to antibiotic treatment by analyzing existing gut metagenomic data from antibiotic treated individuals^49–53^ (Table S1A-C). We first analyzed a study of 12 healthy volunteers who were treated with a 4-day course of combination antibiotics (meropenem, vancomycin and gentamycin)^50^. The gut microbiomes of four of these subjects harbored pMMCAT prior to treatment. In three of these, the plasmid initially became undetectable (when Bacteroidales populations dropped dramatically^50^) and then reappeared by the 42-day time point, when the Bacteroidales abundances had normalized. In contrast, in one subject who initially carried pMMCAT, the plasmid was lost and never reappeared, suggesting that the pMMCAT-harboring strain(s) were eradicated. Two volunteers who did not originally have the plasmid acquired it by day 42, suggesting their gut microbiota became colonized by a new pMMCAT-harboring strain(s) after a niche opened during the antibiotic treatment or the relative abundance of pMMCAT increased above the level of detection. Likewise, we analyzed metagenomes from a study of six healthy volunteers in the USA who were subjected to a challenge of enterotoxigenic *E. coli* (ETEC), causing mild to severe inflammatory diarrhea, followed by ciprofloxacin treatment^51,52^. All six volunteers had pMMCAT and retained it after the ETEC and antibiotic challenge when the microbial community normalized. Finally, we analyzed two cohorts of fecal microbiota transplants (FMT) from healthy donors to 17 patients with Crohn’s disease^53^ or recurrent *C. difficile* infection^49^. Of the five patients that had pMMCAT prior to the FMT, all except one retained it. Of the twelve patients that did not carry pMMCAT prior to the FMT, most acquired it later either from the FMT donor (eight) or from another source (one). While circumstantial, these results suggest that bacteria harboring pMMCAT may be mostly retained during a perturbation, whereas they may be more readily acquired if a new niche becomes available, possibly explaining why this plasmid has become so prevalent, especially in industrialized countries.

### Transcriptional profiles in pMMCAT biofilms

To gain more insight into pMMCAT-mediated biofilms in PhvCL11, we compared gene expression profiles in biofilms grown under static conditions and in fecal pellets from monocolonized mice, prior to and after metronidazole treatment (Table S2, Fig 5). The *eps* genes are expressed at low levels in mice prior to treatment but are strongly induced by metronidazole treatment (Fig. 5A-B), consistent with the MiPACT observations. The *mfa* genes are moderately expressed prior to treatment and two of them are upregulated by metronidazole. Therefore, in the fecal pellet, Mfa-mediated aggregation may also occur outside of the mucus layer but these cells would not be covered with an extracellular polysaccharide matrix that is a hallmark of biofilms. Both loci are also highly expressed in the *in vitro* biofilms under static conditions.

**Figure 5.**
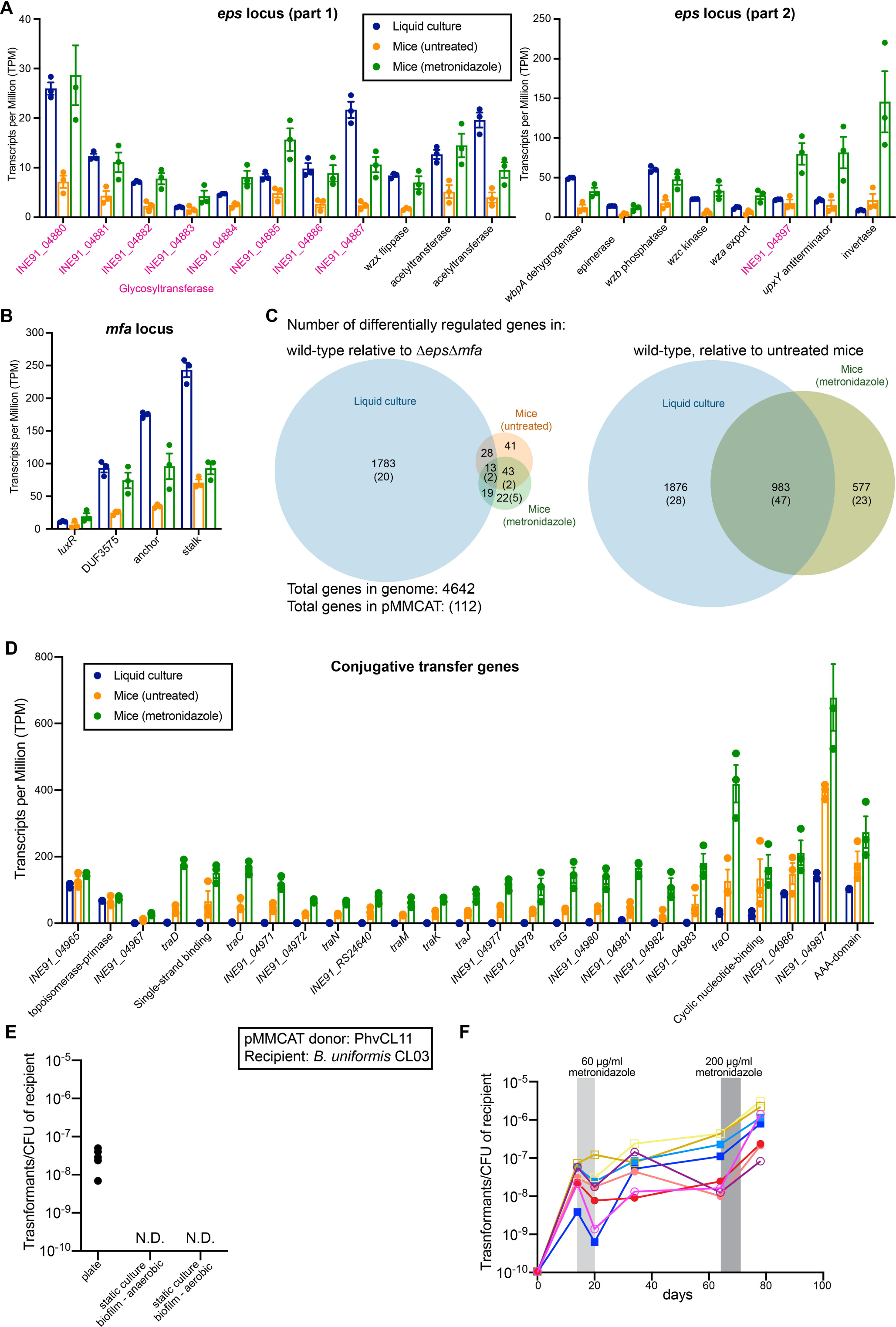
Transcriptional profile of PhvCL11 pMMCAT biofilms. (3 mice per condition). **A-B.** Expression levels (transcripts per million, TPM) of the two main operons of the *eps* locus (A) and the *mfa* locus (B) of PvCL11. **C.** Venn diagrams of the numbers of differentially regulated genes in the comparisons listed. Genes in pMMCAT are shown in parentheses (excluding genes of the *mfa* and *eps* loci for the *ΔepsΔmfa* comparison). **D.** Expression levels (TPM) of the genes involved in conjugative transfer of pMMCAT. **E.** Transfer frequency of pMMCAT from PhvCL11 to *Bacteroides uniformis* CL03T12C37, mixed at equal ratio on plates or in liquid culture biofilms. Five independent replicates are shown. N.D. = below detection limit. **F.** Transfer frequency of pMMCAT from PhvCL11 to *Bacteroides uniformis* CL03T12C37 in eight gnotobiotic mice. Mice were subjected to two metronidazole pulses (60 µg/ml and 200 µg/ml). Mice were co-housed in pairs and are shown in similar colors and with the same symbol. Four mice of each sex were used.

When comparing gene expression profiles under static conditions of the wild-type PhvCL11 biofilms and the isogenic *ΔepsΔmfa* planktonic cells, approximately 40% of the genome (1865 genes) are differentially regulated (Fig. 5C, Table S2). In contrast, in colonic biofilms, transcription of only 106 genes is differentially regulated, 34 of which are also differentially regulated in static culture biofilms. The MiPACT-HCR imaging shows that biofilm formation is preferentially induced in bacteria near the mucus layer, therefore in bulk RNA-Seq the biofilm-associated gene expression may be obscured by non-biofilm producing cells.

Among the 39 genes that are differentially regulated between the wild-type and *ΔepsΔmfa* in metronidazole-treated but not in untreated mice (Table S2), the most striking difference is a 14-gene locus (INE91_04565-04582) comprising an ATPase potassium pump, a gene encoding a putative chloramphenicol acetyltransferase, and a putative extended spectrum β-lactamase resistance gene. This antibiotic resistance locus is also highly upregulated in static culture biofilms. The two latter genes are not expected to increase resistance to metronidazole, which belongs to a different chemical family and has a killing mechanism unrelated to that of chloramphenicol and the β-lactams^54^. However, their differential regulation suggests that cells in a biofilm undergo a regulatory program promoting protection from chemical insults. Interestingly, another potassium uptake gene (INE91_03212), part of a multidrug efflux locus (INE91_03211-3215), is also upregulated. Potassium fluxes are implicated in metabolic processes in *Bacillus subtilis* biofilms, allowing long-range electrical signaling to recruit *Pseudomonas aeruginosa* to form mixed-species biofilms^55^.

Conversely, there is one region (spread across two contigs) whose expression is reduced in the wild-type relative to the *ΔepsΔmfa* mutant during metronidazole treatment (INE91_RS00005-INE91_00114 and INE91_04844-04869). The region encodes a lysogenic phage whose genes are among the most highly expressed in *ΔepsΔmfa* under metronidazole treatment. This prophage belongs to the LoVEphage (Duplodnaviria; Heunggongvirae; Uroviricota; Caudoviricetes), which were recently discovered in gut metagenomes and are highly prevalent^56^. It is also expressed in the wild-type but to a lesser extent. Metronidazole is thought to kill cells *via* the formation of a nitro-radical anion (R-NO_2_•−) which causes oxidative damage to DNA leading to strand breaks^54^. As the wild type is more protected from metronidazole in a biofilm, it likely undergoes less DNA damage leading to decreased prophage induction.

For pMMCAT genes, the most striking change in expression is the invertase regulating the inversion of the *eps* locus promoters and the six upstream genes (INE91_RS24245, INE91_04901-04905), which encode small hypothetical proteins with no characterized domains. These are all highly upregulated in metronidazole-treated biofilms but not in the *ΔepsΔmfa* strain.

### pMMCAT transfer

The pMMCAT genes involved in conjugative transfer (including genes of the type IV secretion system) are highly upregulated by metronidazole treatment in the gut, suggesting stressors may induce pMMCAT transfer (Fig 5D). Using antibiotic selection markers in the chromosome and in pMMCAT, we quantified the plasmid transfer frequency from PhvCL11 to a strain of *B. uniformis* (BuCL03) isolated from a different person (Fig. 5E). Donor and recipient strains were mixed at equal ratios and spotted together in a plate or allowed to form a biofilm in a 96-well plate. After 24 hours, cells were recovered and plated to quantify the populations of donor, recipient and transconjugants (recipient with pMMCAT). The transfer frequency in plate-grown co-cultured bacteria was approximately 10^-8^ transformants/recipient (Fig 5E). This frequency is one to two orders of magnitude lower than other well-studied mobile genetic elements in *Bacteroides*^57–59^. As expected by the low level of transcription of the conjugation genes during *in vitro* biofilm growth, no transfer was detected in biofilms grown in liquid medium. To evaluate pMMCAT transfer in the colon, germ-free mice were co-colonized with donor and recipient strains for two weeks and subsequently subjected to two rounds of metronidazole pulses of increasing strength, roughly one month apart (Fig. 5F). Prior to metronidazole treatment, the frequency of transconjugants detected was highly variable across mice, sometimes even between co-housed animal pairs. After an initial decrease in CFU due to bacterial death from the first metronidazole pulse, the number of transconjugants increased over the following week for four of the eight mice and stayed mostly stable until the beginning of the second pulse. After the second pulse, there was not a reduction in transconjugant CFUs as detected after the first pulse, rather, all mice showed a ∼1-2.5 log increase in transconjugants, whereas both donor and recipient strain abundances remained relatively constant (Fig. S3). This observation is consistent with the RNA-Seq results, which demonstrate that this elevated concentration of metronidazole induces genes involved in conjugal transfer, however, as there was no decrease in transconjugant levels following this treatment, pMMCAT protection is likely also a contributing factor. In the human gut, pMMCAT-harboring bacteria eventually predominate, suggesting that additional factors may be necessary for pMMCAT to become stably fixed in a recipient population.

### pMMCAT is highly conserved and widely distributed in human gut Bacteroidaceae and Parabacteroides species

To determine the prevalence of pMMCAT across the Bacteroidota phylum, we analyzed 12,456 genomes of non-redundant isolates available from NCBI (Table S3). We used the sequence of pMMCAT from *Bacteroides cellulosilyticus* CL06T03C01 as query with a threshold of 94% coverage in a *blastn* search. pMMCAT was present in the three Bacteroidales genera most common in the human gut of industrialized populations: *Bacteroides*, *Parabacteroides* and *Phocaeicola* (Fig. 6A, Table S3)^1,2,60^. Although pMMCAT was prevalent in most species of these genera, it was largely absent from *B. fragilis*, *B. salyersiae* and *Ph. plebeius*. We also searched for pMMCAT in all sequences available on the blastn web interface of NCBI in the default nr/nt nucleotide collection, confirming its absence in other non-Bacteroidota genera. Interestingly, we did not detect pMMCAT in isolates from non-human primates, pets or livestock (Fig. 6B), even though they contain *Bacteroides*, *Parabacteroides,* and *Phocaeicola*. In contrast, other widespread mobile genetic elements in the gut Bacteroidales are found in isolates from pets and livestock^16,61–63^. The plasmid was frequently found in human-derived strains from the USA, Asia (China, Japan) and some countries in Europe, revealing its prevalence in industrialized countries (Fig. 6C).

**Figure 6.**
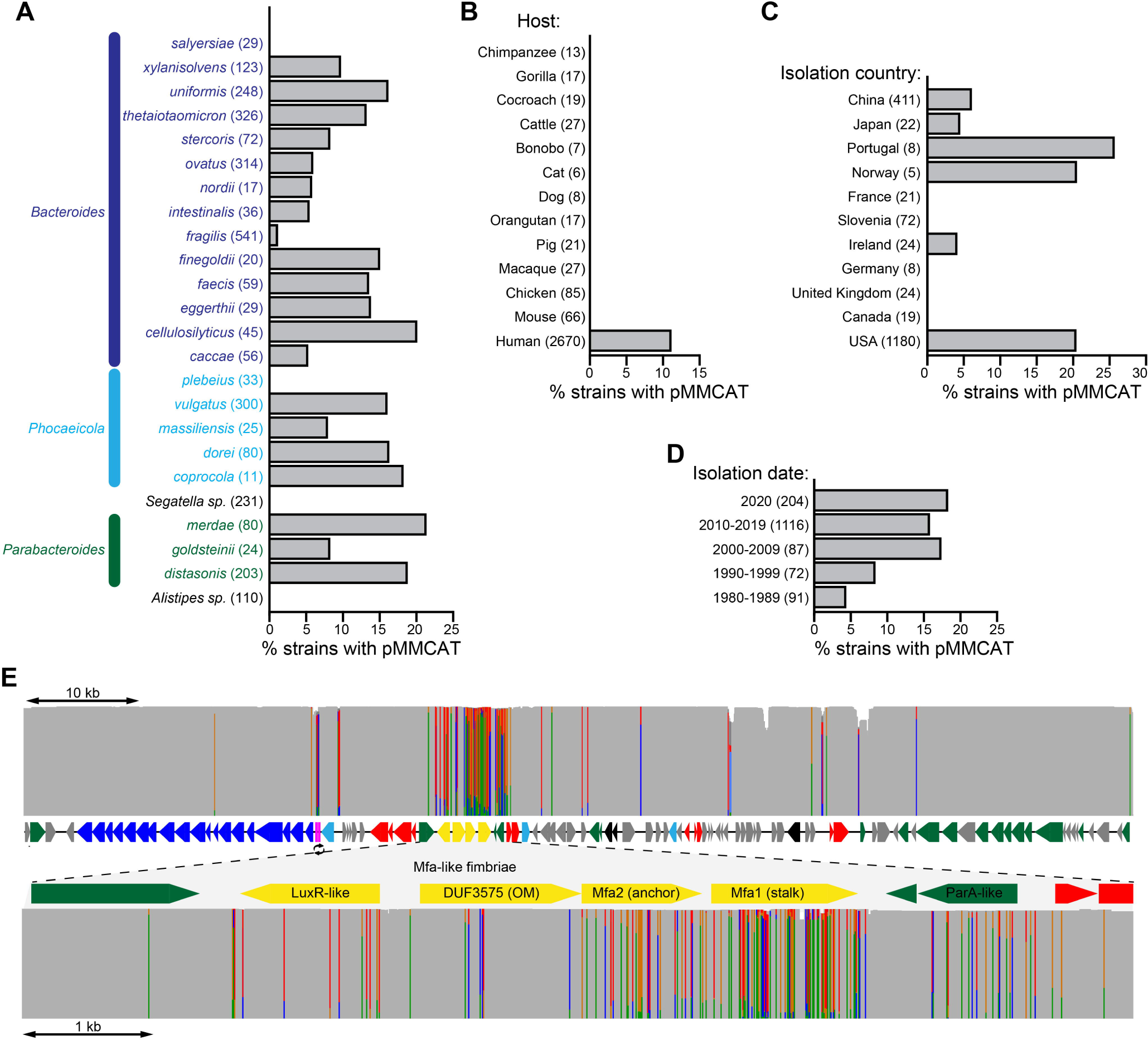
Prevalence of pMMCAT in isolate genomes. **A.** pMMCAT prevalence in Bacteroidales species common in the human gut. Numbers in parentheses indicate the total number of isolates analyzed (non-redundant sequenced genomes). **B.** pMMCAT prevalence in *Bacteroides*, *Parabacteroides*, and *Phocaeicola* by host. Numbers in parentheses indicate the total number of isolates analyzed. **C.** pMMCAT prevalence by country of isolation, for human strains of *Bacteroides*, *Parabacteroides*, and *Phocaeicola*. Species found to not usually harbor pMMCAT were excluded (*B. salyersiae*, *B. fragilis*, and *Ph. Plebeius*). **D.** pMMCAT prevalence by decade of isolation, for human strains of *Bacteroides*, *Parabacteroides*, and *Phocaeicola*. Species that do not typically harbor pMMCAT were excluded (*B. salyersiae*, *B. fragilis*, and *Ph. plebeius*). Slovenian isolates (72 isolates from the 2020s) were also excluded as pMMCAT appears to be absent from this country. **E.** Sequence conservation across pMMCAT (top panel) and the *mfa* locus (bottom), based on a multiple sequence alignment of all 292 pMMCAT sequences from isolate genomes. Grey = no variation at that position compared to the reference. For variable positions in the alignment, the fraction of sequences with each nucleotide is represented by the height of the bar of that color (blue = C, red = T, green = A, brown = G). pMMCAT genes are colored as indicated in Fig. 1A.

We next screened using PCR for the presence of pMMCAT in 64 Bacteroidales clinical strains isolated from 1980-1997 from Boston area hospitals^64^, excluding species that rarely contain pMMCAT (e.g. *B. fragilis*). We also analyzed sequenced genomes of a collection of 100 clinical isolates from this time period from the NIH Clinical Center in Bethesda, MD^65^. These analyses revealed a lower prevalence of pMMCAT in these Bacteroidales strains, especially those isolated prior to 1990, and has increasingly spread in the last two decades (Table S4, Fig 6D). However, as these collections are limited to clinical isolates from two cities, we further analyzed metagenomes from a few available archeological fecal specimens. We analyzed two samples of medieval coprolites/latrine sediments (Latvia and Israel)^66^, the colon contents of the Copper age iceman (Ötzi)^67,68^, pre-Columbian coprolites in America (12 samples from 500-600 A.D., Mexico^69,70^ and Puerto Rico^71^) and Neanderthal coprolites (7 samples from Spain)^72^ (Table S1D). All seven human samples from pre-Columbian Mexico coprolites had reads that mapped to 5-46% of pMMCAT. The regions of pMMCAT with mapped reads were those involved in conjugation and plasmid maintenance, shared among diverse conjugal elements and therefore not pMMCAT-specific (Fig. S4). Notably, two samples had 35-46% of the pMMCAT backbone (lacking *mfa* and *eps* regions), revealing that some of its mobilization components have been circulating in humans for at least 14 centuries. In contrast, all regions of pMMCAT were undetectable in other archeological samples, where the abundance of *Bacteroides*, *Parabacteroides* and *Phocaeicola* is low.

Analysis of the sequences of pMMCAT retrieved from all sequenced isolates shows that it is highly conserved (higher than 93% identity over the length of the plasmid for all samples and higher than 99.97% for most). To evaluate this conservation more systematically, we generated a multiple sequence alignment of all pMMCAT sequences from 292 genomes and evaluated conservation across each nucleotide (Fig. 6E). Indeed, the plasmid is highly conserved across most of its sequence, with a variation hotspot in the *mfa* fimbriae locus, particularly the gene encoding the stalk protein which composes the main structural component of the fimbriae and influences binding specificity^73^. To better study this variation, we reconstructed a maximum-likelihood phylogeny of the plasmid (Fig. S5). This analysis showed that the variation could be explained by the independent emergence of *mfa* locus variants in at least eight independent events, with one main variant which we call pMMCAT-var1. Most SNPs were not shared across variants. Altogether, *mfa* variants represent 37% of sequences. We did not find similar genomic regions that might have given rise to these variants *via* recombination. Intriguingly, isolates from the USA contained not only the predominant *mfa* sequence but also many variants, whereas isolates from China lacked the predominant *mfa* sequence present in the US and only contained variants, primarily pMMCAT-var1. Therefore, these variants might have emerged in China before being transferred to isolates in other countries through migration. We also detected a common variant, derived from pMMCAT-var1, in several isolates from China in which the entire *eps* locus is absent (Table S3C, Fig. S6, in 44 isolates). While various small regions of pMMCAT (notably not the *eps*) were found in many genomes, none were conserved and common except for this *eps*-deficient variant, found exclusively in China. The plasmid phylogeny also reveals 16 additional instances of the likely spread of pMMCAT among species from the same person, in two isolate collections from the USA which were not available at the time we conducted our previously reported analysis^74,75^, once again highlighting how this plasmid is frequently spreading within species in individuals. pMMCAT is >99.995% identical between these co-resident species, which group together in the pMMCAT phylogeny (e.g. from subject DFI.6: *Pa. distasonis* DFI.6.54, *Ph. vulgatus* DFI.6.58, *B. thetaiotaomicron* DFI.6.61 and *B. finegoldii* DFI.6.73).

To determine the extent to which pMMCAT has changed in the last two plus decades, we sequenced the genomes of all isolates from the Boston collection that carried pMMCAT. Six of the eight 1980s-1990s isolates with pMMCAT (three from the Baltimore collection and three from Boston) were 99.99% identical to the predominant sequence with 3-9 SNPs over the entire 98 kb plasmid (Fig S5). This striking sequence conservation relative to the current dominant variant suggests that pMMCAT has remained mostly the same as it spread widely in human populations. Interestingly, one isolate from the 1990s (TS50) had an *mfa* variant divergent from any recent variants, suggesting that *mfa* diversification has been ongoing since the 1990s in a small proportion of the population in the USA, but has not become widespread.

### pMMCAT is globally ubiquitous

Since available sequenced genomes are limited to a few countries, we used publicly available metagenomic datasets of 4467 human fecal metagenomes from healthy adults in 26 countries to quantify the prevalence of pMMCAT across global human populations. Using a threshold of 90% coverage, we found pMMCAT present in 24 of the 26 countries (Fig. 7A, Fig. S7A, Table S1E). More than half of study subjects in Canada, USA, Brazil, and Thailand carried pMMCAT, while prevalence was greater than 20% in several countries from western Europe, Asia, Africa, Australia, and Israel. pMMCAT was not detected in the Peruvian population and only detected in a few subjects from Fiji^15^ and Madagascar^76^. Moreover, pMMCAT was also absent from hunter-gatherers in Tanzania^77^ but present in an agricultural population from this country^68^. Taxonomic analysis of the microbial composition of the microbiota from these generally non-industrial populations indicated that the Bacteroidales are largely of the genus *Segatella*, whereas the genera that normally harbor pMMCAT are rarely present (Table S5A). Accordingly, all pMMCAT-positive metagenomes were found to contain one or more species of *Bacteroides*, *Phocaeicola*, or *Parabacteroides* (Table S5A). Surprisingly, pMMCAT prevalence was very low in India and Bangladesh despite the fact that 67 out of 105 subjects in India had communities dominated by *Bacteroides* and *Phocaeicola*, including many of the host species with high prevalence of pMMCAT (Table S5B). To gain further insights into the geographic prevalence of pMMCAT, we analyzed 950 global sewage metagenomic samples from 101 countries^78^ (Fig. 7B, Table S1F, Table S6). Strikingly, the plasmid was detected in sewage from 56 countries from all continents except Antarctica, highlighting its global distribution.

**Figure 7.**
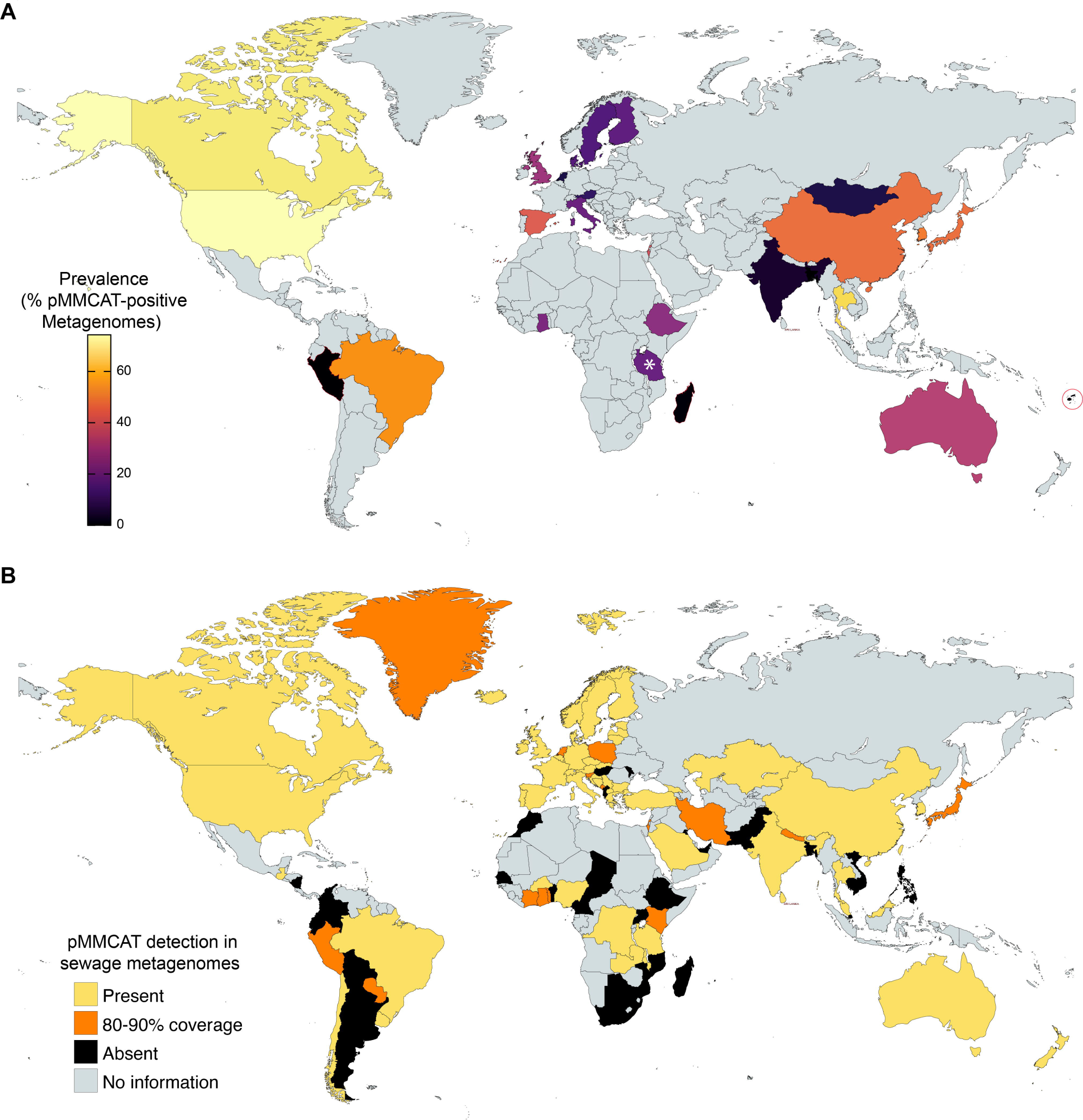
pMMCAT is globally distributed. **A.** Prevalence of pMMCAT in global gut metagenomes. The white asterisk indicates a second cohort in Tanzania (Hadza hunter-gatherers) with 0% prevalence. **B.** pMMCAT prevalence in global sewage metagenomes. A country is shaded as yellow if pMMCAT was detected (>90% sequence coverage) in at least one sewage sample. Countries where only partial coverage of pMMCAT was detected (80-90% of the sequence) are shaded in orange. Number of sewage metagenome samples per country is shown in table S6.

To determine if pMMCAT carriage is influenced by age, we evaluated infant and children gut metagenomic datasets from ten countries, showing that pMMCAT could be detected from infancy (Table S1E-F) and the prevalence in children was comparable to that in adults (Fig. S7B). We also sought to determine the prevalence of pMMCAT in gut metagenomes of patients with various intestinal disease states. We found no consistent difference between healthy adults and subjects with colorectal cancer (Japan^79^, Fig. S6C). pMMCAT prevalence in individuals with Inflammatory Bowel Disease (IBD) appeared lower than in healthy controls (Fig S7D, Table S1G), in cohorts from Spain^80^, Netherlands^81^ and the USA^81–83^. Interestingly, for individuals where pMMCAT was present, its abundance (proportion of total reads mapped to pMMCAT) appeared higher in IBD patients relative to healthy subjects (Fig. S2E), suggesting that in many subjects there is either increased copy number or increased abundance of the strains harboring the plasmid.

We next determined if pMMCAT is present in other habitats or hosts. pMMCAT is absent in metagenomes of other human body sites (Table S1H) (2971 samples from oral, nasopharyngeal, lung, skin, and vaginal microbiomes) with only very rare detection in two skin samples (2 subjects out of 1530) and three oral samples (5 out of 879). Since analysis of metagenomes provides much greater sensitivity than scrutinizing isolate genomes, we looked for a potential animal or environmental reservoir for pMMCAT in a vast array of published metagenomic datasets. pMMCAT was absent from non-human primates, cats, dogs, and wildlife (4877 animal gut metagenomes were analyzed) (Table S1I). It was very occasionally present in livestock and in animals in captivity (four Golden snub-nosed monkeys, 24 lab mice, one goat, one pig, two cows, and eight chickens). Given the low prevalence, we hypothesize that these animals harbor Bacteroidales strains acquired from a human source. Similarly, pMMCAT was absent in 10,044 global non-sewage environmental samples analyzed (marine environments, plants, soil, built environments) (Table S1J), except in seven seawater/sediment/seawall biofilm samples where human fecal contamination is likely. Therefore, pMMCAT is highly specific to the human intestinal microbiome. In a survey of hospital surfaces in Singapore^84^ (Table S1J), pMMCAT was detected in high-touch surfaces such as bedside lockers (3), a pulse oximeter and a bed rail, likely due to fecal contamination, highlighting a potential route for person to person spread.

## DISCUSSION

The modern human lifestyle has rapidly changed our gut microbiome and its ecology. As such, compensatory mechanisms can arise that confer better fitness under these new conditions. Indeed, the large conjugative plasmid pMMCAT, now ubiquitous in diverse human populations, appears to have undergone recent spread, with evidence that it continues to spread, and is still rare in some populations (primarily non-industrialized). Despite great interest in horizontal gene transfer in the gut microbiome, there have been few efforts to experimentally identify their evoked phenotypes and connect these functions with the ecological and evolutionary dynamics of the corresponding mobile genetic elements. While most MGEs are highly variable, subject to frequent recombination, and geographically clustered^15,17^, a few “fitness optimal” combinations are conserved. To our knowledge, only one other 2.7-kb plasmid is as highly prevalent as pMMCAT, but this very small cryptic plasmid appears to be parasitic on its host cell^57^. Our data show that pMMCAT confers a significant fitness advantage to the strains that acquire it, potentially explaining its rapid fixation in human populations.

Here, we present evidence that pMMCAT promotes the formation of a multispecies biofilm in the colon, especially at the mucus layer, and is likely to impact ecosystem dynamics. Cooperative phenotypes such as biofilms and their role in ecosystem properties are poorly characterized in the diverse intestinal community. To our knowledge, there are very few reported cases of cooperation mediated by shared mobile genetic elements^85–89^. In addition, the amount and stability of cooperative interactions among species in the intestinal microbiota remains an active topic of discussion^90,91^. Notably, one example of a conjugative plasmid promoting bacterial biofilm development was discovered in *E. coli,* where expression of the conjugation machinery from the F plasmid leads to induction of chromosomal genes that mediate biofilm formation *in vitro*^92^. Other plasmids have also been shown to increase the amount of biofilm formation *in vitro* in *E. coli* through so far unknown mechanisms^93,94^.

Gut mucosal biofilms have been studied in pathogens and in colorectal cancer^29,46,95^. In contrast, biofilms formed by the microbiota of the healthy gut remain undercharacterized^95^. In Bacteroidales species, biofilms have only been studied *in vitro*^30,31,37,38,96,97^. MiPACT-HCR enabled us to study expression of genes involved in the production of the EPS biofilm matrix in the colon with unprecedented 3D spatial resolution. While HCR is significantly more sensitive than conventional FISH^98^, it may only reliably detect very highly expressed genes in bacteria, especially in whole tissue samples^43,44^. Therefore, we cannot rule out that *eps* is also expressed at lower levels further away from the mucus layer. However, our data support that the strongest *eps* biofilm matrix expressing bacterial cells are located at the mucus layer and on some plant material in the colon. These are very different from the mucosa-attached pathogenic and colorectal cancer biofilms, which are in direct contact with the host tissue^28,29,95^. The type of biofilms formed by pMMCAT-harboring strains, their matrix properties, metabolic interactions, and exact regulatory programs are likely highly dependent on which strains are present in the community. Several factors determine which bacteria can partake in multispecies biofilms. For example, production of specific surface attachment proteins may allow bacteria to form or join an established biofilm. In contrast, antagonism can lead to exclusion of unwanted competitors and cheaters^26,99^. Moreover, one exciting possibility is that, if pMMCAT-mediated biofilms similarly increase survival outside the host, this plasmid could facilitate the co-transfer of strains from person to person.

It will be interesting to study pMMCAT’s evolution in the future, especially in the countries where fimbriae variants (or *mfa*-deficient variants) are common. We speculate that some of this important variation may arise in geographic clusters due to selective pressures such as diet, with new variants spreading in other countries due to globalization. As much of this variation alters the fimbrial stalk protein and may alter binding specificity, it will also be important to evaluate its impact in community dynamics.

Finally, we provide evidence that pMMCAT transfer is induced by metronidazole treatment. Therefore, under this stress condition (and likely others identified *in vitro*), not only are pMMCAT-harboring cells less vulnerable, but also promotes further spread of the plasmid to other bacteria in the community, potentiating community resilience.

## ACKNOWLEDGEMENTS

We thank Paula Montero Llopis of the MicRoN Imaging facility at Harvard Medical School and Otto Cordero for helpful discussions and support for L.G.B.’s training. Confocal microscopy was performed in the Integrated Light Microscopy Core at University of Chicago, which receives financial support from the Cancer Center Support Grant (P30CA014599). We are thankful to Christine Labno for technical assistance with confocal microscopy (RRID: SCR_019197). Gnotobiotic mouse experiments were performed in the Gnotobiotic Research Animal Facility of the University of Chicago, were we received guidance from Betty Theriaut and assistance with breeding from Kimberly Campbell, Melanie Spedale and Kristin Kolar. We thank Tatyana Golovkina, Sam Light and Eric Pamer for germ-free mice. We are grateful to Anitha Sundararajan, Rita Smith, Carolyn Metcalfe, Che Woodson, and Victoria Burgo of the DFI Microbiome Metagenomics Facility for RNA sequencing and whole genome sequencing, and to the DFI Bioinformatics core team, especially Ramanujam Ramaswamy and Huaiying (Eddi) Lin, for analyses and helpful discussions. We thank Emily Fogarty for helpful discussions and for sharing several metagenomic datasets, Sam Light, Mark Mimee, and members of the Comstock and Waldor labs for useful feedback. We are grateful to Mary Delaney, Lynn Bry and Andrew Onderdonk for sharing the historical Bacteroidales isolate collection. We thank Jennifer Teschler for technical guidance with MiPACT-HCR protocols. This work was funded by K99AI167064 to L.G.B and R01AI093771 to L.E.C., from the NIH/National Institute of Allergy and Infectious Diseases and by the Duchossois Family Institute. N.M.E. is supported by NIH T32 GM144292, M.K.W. is an Investigator of the Howard Hughes Medical Institute. The funders had no role in study design, data collection and interpretation, or the decision to submit the work for publication.

## AUTHOR CONTRIBUTIONS

L.G.-B. and L.E.C. designed the studies and wrote the manuscript. M.J.C. edited the manuscript. L.G.-B, N.S., and K.F. performed experiments. L.G.-B, N.M.E., M.L.S., K.H. and K.F. assisted with gnotobiotic mouse work. L.G.-B., M.J.C and A.G.C. performed computational analyses. F.H.Y., A.T.K. and M.K.W. provided intellectual expertise and technical guidance. All authors read and commented on the manuscript.

## FIGURE LEGENDS

**Figure S1. pMMCAT is circular and present at about one to three copies per genome. A-C.** Genome assembly graphs from three representative sequenced genomes with pMMCAT. All plasmids are shown with their predicted average copy number. **D-E**: GroEL immunoblots as loading controls for immunoblots shown in Figure 1B-C. **F.** Immunoblot of EPS present in the supernatant of strains of PhvCL11 showing a high molecular weight polysaccharide.

**Figure S2. MiPACT-HCR 3D imaging of biofilms in the colon. A.** Mouse colon embedded in hydrogel, shown prior to and after clearing. **B.** Region of the *eps* locus where the HCR probes anneal. **C-D.** Mice were monocolonized with PhvCL11 (constitutively expressing *mCherry*) and colons were imaged prior to **(C)** and during metronidazole treatment **(D)**. For each set of images, the top row shows the 3D Blend Projection in Imaris, and the bottom row shows a representative slice along the z-stack. DAPI (shown in blue) was used to stain DNA (host cell nuclei and bacterial cells), while autofluorescence of fecal pellet debris was imaged in the far red (shown in cyan). Debris are known to exhibit some degree of autofluorescence across most wavelengths. HCR probes against *mCherry* (shown in red) were used to detect all transcriptionally active PhvCL11 cells, while probes against the *eps* locus (shown in green) were used as a proxy for expression of biofilm genes. White arrows indicate examples of cells expressing *eps* inside plant debris compartments.

**Figure S3.** pMMCAT PhvCL11 donor (**A**) and BuCL03 recipient (**B**) abundances in mouse transfer experiments.

**Figure S4. Coverage of reads recruited to pMMCAT in coprolites.** Metagenomes obtained from the same sample using different DNA extraction methods are indicated by a black bar on the right of the coverage plots (e.g. sample Zape 5A was analyzed comparing 5 different DNA extraction methods).

**Figure S5.** Maximum likelihood tree of complete pMMCAT sequences from isolate genomes. Magenta branches = *mfa* sequence variants. Purple box = main *mfa* variant. Blue = isolates from China. Red = isolates from prior to 2000. Dashed line box illustrates an example of multispecies transfer within one person’s microbiota (individual DFI.6).

**Figure S6**. Alignment of *eps*-deficient variants to the pMMCAT reference sequence at the bottom (from *B. cellulosyliticus* CL06T03C01). For each aligned sequence, a maroon ribbon indicate regions where the sequences align. Inverted triangles indicate an insertion of several nucleotides in the alignment, whereas a maroon tick above the ribbon indicates insertion of a single nucleotide. Mismatches are indicated by white ticks (or spans of white ticks, such as in the *mfa* region) and larger deletions (such as the complete *eps* locus) are shown as white rectangles. pMMCAT genes in the reference are colored as indicated in Fig. 1A.

**Figure S7. A.** Prevalence of pMMCAT in global microbial gut metagenomes. Numbers in parenthesis indicate the number of genomes analyzed. **B.** pMMCAT prevalence in children gut microbiome metagenomes as compared to adults. **C.** pMMCAT prevalence in metagenomes from colorectal cancer patients as compared to healthy individuals. **D.** pMMCAT prevalence in metagenomes from IBD patients as compared to healthy individuals. **E.** Histograms of the percent of metagenomic reads that map to pMMCAT in IBD patients as compared to healthy controls.

**Table S1. Read recruitment analysis to pMMCAT in human, animal and environmental metagenomic datasets.**

**Table S2. RNA-Seq analysis of PhvCL11 wild-type and ΔepsΔmfa in static cultures and in monocolonized mice. S2A.** Mapped read counts per sample. *S2B.* Fold change and probability values for each gene according to DESeq2 and EdgeR. *S2C.* Transcripts per million (TPM) for each gene and condition.

**Table S3. Similarity search (blastn) of pMMCAT in sequenced genomes from isolates. S3A.** Presence or absence of pMMCAT in each genome, as well as relevant sample metadata. *S3B.* Redundancy analysis to identify nearly isogenic genomes where multiple isolates from the same species where isolated from the same subject (oner per subject was kept). *S3C.* Genomes harboring the pMMCAT variant without the *eps* locus, as well as relevant sample metadata.

**Table S4. pMMCAT prevalence in strains isolated between 1980-1999 in Boston area hospitals.**

**Table S5. Species abundances in metagenomes, estimated using Kraken.** Each lane represents one metagenome sample (metadata available in Table S1). Total number of fragments classified to each species in each sample is reported after the species name in the row.

**Table S6. pMMCAT detection in sewage metagenomes.** Table S7. Primers (S7A), plasmids (S7B), synthetic DNA fragments (S7C) and bacterial strains (S7D) used in this study.

**Video S1. Strains harboring pMMCAT form tight aggregates in liquid culture.** Part 1: pure culture of PhvCL11 wild-type or *ΔepsΔmfa.* Part 2: Co-culture of the five indicated strains (all harboring pMMCAT) isolated from the same person (CL11), mixed at equal OD in mid-exponential phase.

**Video S2. MiPACT-HCR 3D imaging of the colon of mice monocolonized with PhvCL11 in the absence of metronidazole treatment.** A strain of PhvCL11 constitutively expressing *mCherry* was used. DAPI (shown in blue) was used to stain host cell nuclei and bacterial cells, while autofluorescence of fecal pellet debris was imaged in the far red (shown in cyan). HCR probes against *mCherry* (shown in red) were used to detect all transcriptionally active *Ph.v.*CL11 cells, while probes against the *eps* locus (shown in green) were used as a proxy for expression of biofilm genes.

**Video S3. MiPACT-HCR 3D imaging of the colon of mice monocolonized with PhvCL11 with metronidazole treatment.** Legend as for Video S2.

**Video S4. MiPACT-HCR 3D imaging of the colon of mice monocolonized with PhvCL11 Δ*epsΔmfa* with metronidazole treatment.** Legend as for Video S3.

## MATERIALS AND METHODS

### Media and growth conditions

*E. coli* S17-1 λ*pir*^100^ was grown aerobically at 37°C in lysogeny broth (LB) medium. Bacteroidales strains were grown at 37°C under anaerobic conditions (10.0% H_2_, 10.0% CO_2_, balance N_2_) in supplemented basal medium^101^ (liquid cultures) or brain heart infusion plates supplemented with 5 mg/liter hemin and 2.5 μg/liter vitamin K1 (BHIS)^101^. Antibiotics were used at the following concentrations, where appropriate: 100 μg/ml carbenicillin, 5 μg/ml erythromycin, 200 μg/ml gentamicin, 6 μg/ml tetracycline, 100 ng/ml anhydrotetracycline.

### Plasmid and strain construction

All primers, synthetic DNA blocks, plasmids and strains used in this study are listed in tables S7A-D, respectively. PCR reactions were carried out using Phusion high-fidelity DNA polymerase (New England Biolabs, Ipswich, MA). Plasmids were digested using NEB high-fidelity restriction endonucleases in Cutsmart buffer and plasmid assembly reactions were performed with NEBuilder HiFi DNA assembly, following manufacturer’s directions. All constructs were verified using Oxford Nanopore Technologies sequencing at Plasmidsaurus (Eugene, OR) or Primordium (Louisville, KY). Bacteroides conjugation and counterselection for mutant construction was performed as described previously^101^. Briefly, the plasmid was transferred via conjugation from *E. coli* to the target Bacteroidales strain and transconjugants were selected using the appropriate antibiotic and gentamycin. Transconjugants were grown for 4-8 hours in supplemented basal medium and plated on BHIS with 100 ng/ml anhydrotetracycline for counterselection. Mutants were verified by loss of the plasmid selection marker and by PCR (Table S7A). Since pMMCAT is present at 1-3 copies per cell, care was taken to verify that all selected clones harbored only pMMCAT with the desired mutation. For each mutant, three independent clones were tested in phenotypic assays.

### EPS and Mfa antibody generation and immunoblotting

Custom antibodies against the Mfa stalk protein were raised as described previously^102^. Briefly, INE91_04914 from PhvCL11 was cloned into pET16b (MilliporeSigma, Darmstadt, Germany) for construction of an N-terminal His-tagged fusion protein for expression in *E. coli* BL21(DE3)^103^. Following induction with IPTG, the His-tagged protein was purified using the ProBond Purification System (Thermo Fisher Scientific) and the eluted protein was dialyzed against PBS. This purified protein was used to raise antiserum in rabbits using the Express-line polyclonal antiserum protocol (Lampire Biologicals). Custom antibodies against the EPS were raised as described previously^104^. Biofilms of PhvCL11 with the *eps* promoters orientation locked “on”-“on” were grown in culture and fixed in neutral buffered formalin. 10^8^ cells per dose were used as the immunogen for generation of rabbit antiserum at Lampire Biologicals, using their Express-Line protocol. This antiserum was adsorbed in two successive rounds with PhvCL11 Δ*eps* cells to remove the antibodies to all surface molecules except the EPS and filter-sterilized. The use of rabbits for antiserum generation was approved by the Institutional Animal Care and Use Committee (IACUC) and the University of Chicago. Polyclonal GroEL antibody (*E. coli*) was purchased from Enzo Life Sciences. Immunoblots were performed as previously described^58^, with a 1:30 dilution of the EPS antibody, 1:100 dilution of the Mfa antibody and 1:1000 dilution for GroEL.

### Crystal violet quantification of biofilms

Strains were grown overnight on supplemented basal medium, diluted to an OD_600_ of 0.1 in pre-reduced medium and allowed to grow until an OD_600_ of approximately 0.5. All cultures were matched to the same OD_600_ and 1.5 mL was added to each well of a 24-well flat-bottom tissue culture treated plate. Four replicates from independent cultures were performed per strain per treatment. After 24 hours incubation, the culture supernatant was carefully removed with a pipette and the biofilm was quantified as previously described with some modifications^37,105^. Briefly, the plate was dried for two hours at 60°C then submerged right side up in distilled water, which was then carefully decanted by inversion of the plate. The plate was briefly dried at 37°C to remove any remaining water and 650 µL of a 1% crystal violet solution in water was added. After 5 hours at room temperature, the solution was discarded by inversion, the plate was washed three times with distilled water and air-dried as before. Crystal violet remaining in each well was resuspended in 500 µL of a 1:4 acetone:ethanol solution. 100 µl of this resuspension was placed in a 96-well plate and its absorbance at 575 nm was quantified in a BioTek Epoch 2 microplate spectrophotometer. If absorbance values were above 1.5, the sample was diluted to preserve linearity and multiplied by the corresponding dilution factor.

### Confocal imaging of liquid culture biofilms

Strains were grown overnight on supplemented basal medium, diluted to an OD_600_ of 0.1 and allowed to grow until an OD_600_ of about 0.2 was achieved. GFP and mCherry cultures of the strains to be tested were matched to the same OD_600_ and mixed with 3 ml of each in a culture tube. After 24 hours anaerobic incubation at 37°C, the culture supernatant was removed using a pipette and the remaining cells were passaged to a 35 mm glass bottom imaging dish (Cellvis). Dishes were exposed to room air for five minutes to allow maturation of the fluorophores inside the biofilm, the biofilm was coated with 250 µl of ProLong Diamond Antifade Mountant (Life Technologies) and covered with a 22 mm × 22 mm coverslip. Samples were imaged on a Leica SP8 3D STED laser scanning confocal inverted microscope using a HC PL APO 40×/1,30 OIL CS2 objective (Leica Microsystems), using LAS X Software (Leica Microsystems). Signal from the different channels was acquired sequentially, collecting all channels in each focus plane, with detectors set to 12-bit mode. GFP signal was collected using a 4881nm argon laser line for excitation and a HyD SMD detector with 500-530 nm window. Signal from mCherry was collected using a 5611nm laser line and a HyD SMD detector with 592-634 nm window. z-stack volumes were set by finding the bottom of the sample just above the dish bottom in the GFP channel and 3 µm z-steps were imaged up to approximately 200 µm total depth. 512 x 512 pixels fields of view were acquired in 4 x 4 mosaic with 10% overlap for each sample. For each strain combination, a minimum of 4 independent replicates were tested. Image visualization was performed with Bitplane Imaris software v. 9.1.2 (Andor Technology PLC), using 3D View with the Blend projection. Image segmentation was carried out in individual fields of view in BiofilmQ using default parameters. After segmentation, thresholds for GFP and mCherry signals were determined by plotting both channels and determining the mean of the signal in the wrong channel. We discarded any cells that had both GFP and mCherry signals above the threshold and whose signal on one channel was less than 1.5 times the signal in the other channel. Plotting of GFP/cell ratios per sample and statistical analyses were done in Prism 10 (GraphPad). Statistical comparison of the mean across samples was performed using one-way ANOVA with Tukey correction for multiple comparisons.

### Gnotobiotic mouse experiments

Gnotobiotic mouse experiments were carried out at the Gnotobiotic Research Animal Facility (GRAF) at the University of Chicago and were approved by the University’s Institutional Animal Care and Use Committee (IACUC). Germ-free C57BL/6 mice were gavaged with 200 µL of an early stationary phase culture of the desired strain or strain combination and were maintained in individually ventilated cages with LabDiet® JL Rat and Mouse/Auto 6F 5K67. All manipulations were made under sterile conditions. For RNA-Seq, metronidazole survival, and plasmid transfer experiments, mice were colonized at 6-10 weeks. For MiPACT-HCR, mice were colonized at 4 weeks. For plasmid transfer experiments, mice were housed in pairs, otherwise they were housed in groups of 3. After two weeks, mice were moved to a new cage with metronidazole in the desired concentration (65-300 µg/ml) with 2% sucrose in the drinking water. After one week, they were returned to regular drinking water. At each time point, fecal pellets were collected under sterile conditions, homogenized in basal medium, serially diluted and plated for CFU counts on BHIS plates with the relevant antibiotics. For RNA-Seq, mice were housed in groups of three and after two weeks of colonization, their drinking water was replaced with 150 µg/ml metronidazole with 2% sucrose. Fecal pellets were collected prior to starting the antibiotic treatment and seven days after the start of metronidazole treatment. Pellets were immediately frozen in an ethanol-dry ice bath and stored at -80°C until further processing.

### 3D Imaging of colon biofilms

MiPACT-HCR imaging of colon biofilms was performed using modifications to the protocol established by Gallego-Hernandez, *et al*.^44^. Hybridization Chain Reaction probes and amplification hairpins were purchased from Molecular Instruments. At each point, three mice per treatment were euthanized, the colons were dissected from the anus to the small intestinal junction, including the cecum and a plug of 0.5% low melt agarose in PBS was added at each end. Samples were fixed in 50 ml methacarn for 48 hours with gentle shaking at room temperature, washed twice for 30 minutes in 70% ethanol, and embedded with 50 ml B4P1 (4% bis-acrylamide (29:1), 1% paraformaldehyde, 0.25% VA-044 in PBS) for 4 days at 4°C. The cecum was removed and the colon was threaded through an RNAse-free 6 mm wide and 90 mm long rubber tube and placed inside a 100 mm long glass culture tube. Inside the anaerobic chamber, B4P1 was bubbled with anaerobic gas mixture and added to fill the culture tube. After 12-16 hours, the samples were retrieved, excess hydrogel on the outside was removed, the colon was placed in 50 ml 8% SDS in PBS pH 8.5 for 5 days in a conical tube, and incubated at 37°C. This incubation step and all subsequent steps were performed with gentle shaking. Samples were washed twice for 30 minutes in 50 ml PBS at 37°C, then were incubated for 16 hours in 10 ml of 600 µg/ml proteinase K in 10 mM Tris pH 7.6 at 37°C. Samples were then placed again in 8% SDS in PBS pH 8.5 for 2 days and washed twice with PBS as before. Cleared colon samples were cut in 3-4 cm long sections and placed in 2 ml hybridization buffer (250 μL/ml formamide, 100 mg/ml dextran sulfate, 2× saline-sodium citrate buffer SSC) with 30 nM of each hybridization probe for three days at 46°C. After an 8-hour wash in 50 mL of 84 mM salt buffer (84 mM NaCl, 20 mM Tris-HCl pH 7.6, 5 mM EDTA pH 7.2, 0.01% SDS) at 52°C, samples were moved to 2 ml of the amplification buffer with amplification hairpins and covered in the dark at room temperature for 3 days. To prepare the amplification buffer with hairpins, 20 µL of each hairpin was incubated in a thermocycler for 90 seconds at 95°C followed by 24°C for 30 minutes, then added to the amplification buffer (100 mg/ml dextran sulfate, 2X SSC buffer). After an 8-hour wash in 50 mL of 337.5 mM salt buffer (337.5 mM NaCl, 20 mM Tris-HCl pH 7.6, 5 mM EDTA pH 7.2, 0.01% SDS) at 48°C in the dark, each sample was incubated for 16 hours in 2 mL PBS with 10 ug/mL DAPI at room temperature in the dark, then moved to RIMS^106^ (40 g of Histodenz in 30 ml of 0.02 M phosphate buffer with 0.1% Tween-20 and 0.01% sodium azide, pH 7.5) with 1 ug/mL DAPI for 16 hours. For imaging, samples were placed on slides with two stacked 1 mm deep press-to-seal silicone isolator sheets. Wells were filled with RIMS, covered with a coverslip and sealed.

Samples were imaged on the LEICA SP8 laser scanning confocal, with the 40× objective and the detectors set to 16-bit mode. DAPI signal (cells/DNA) was detected using a 405 nm laser line for excitation and a photomultiplier tube set to 415-463 nm for detection. AF488 (*eps* HCR probes) was collected using a 488 nm laser line for excitation and a HyD SMD detector with 500-530 nm window. AF546 signal (*mCherry* HCR probes, PhvCL11 cells) was detected using a 539 laser line and HyD SMD detector set to 532-634 nm. Autofluorescence of food particles was detected using a 405 nm laser line and a HyD detector set to 660 – 779 nm. For each sample, a 5 × 5 mosaic (512 × 512 pixels per field of view) was collected (10% overlap with statistical blending), with Z-stacks starting at the coverslip surface to a depth of approximately 220 µm with a Z-step of 3 µm. For each frame and focus plane, 8 line scans were collected and averaged. Each frame was captured along the z-stack and each channel was captured sequentially in each focal plane. Image visualization was done in Bitplane Imaris software v. 9.1.2 (Andor Technology PLC), using 3D View with the Blend projection.

### Quantification of species abundances in co-culture biofilms

Each strain was grown separately overnight in liquid culture as described previously, diluted in pre-reduced medium and allowed to grow to an OD_600_ of approximately 0.5. All cultures were matched to the same OD_600_ and 1 ml of each strain was mixed. After 24 hours of incubation, 2 ml of the planktonic fraction were collected with a pipette into an Eppendorf tube. For the biofilm fraction, all the culture liquid was carefully removed with a pipette and the biofilm was collected into an Eppendorf tube for centrifugation. Samples were centrifuged at 15000 rcf for 2 minutes, the supernatant was discarded and the pellet was stored at -20°C for DNA extraction. Species abundance was quantified at the Duchossois Family Institute Microbiome Metagenomics Facility (DFI-MMF). In brief, DNA extraction was done using the QIAamp PowerFecal Pro DNA kit (Qiagen), followed by fragmentation and library preparation using the QIAseq FX DNA library kit (Qiagen). Libraries were sequenced in an Illumina NextSeq 1000 platform using 2 × 150 reads, producing around 8-10 million paired-end reads per sample. Adaptor trimming and quality filtering was done using Trimmomatic (v.0.39)^107^, then taxonomy profiles were calculated using metaphlan4 (v.4.0.2)^108^.

### Quantification of survival to antimicrobial agents

LL-37 and Bd-A peptides were synthesized by Biotech Peptides and Lifetein, respectively, resuspended in water and filter-sterilized. Liquid culture biofilms were grown as described previously, in 96-well TC-treated plates. Three hours after the biofilm pellets were first visible, the antimicrobial agent was added to the well at the desired concentration, followed by gentle shaking of the plate for 30 seconds. After 24 hours, the contents of each well were homogenized, serially diluted, and spread on BHIS plates to quantify CFUs. Prism 10 (Graphpad) was used to compare the means between strains using unpaired t-tests for each concentration of stressor, with the Holm-Šídák correction for multiple testing. Survival to a metronidazole pulse in mice was carried out as described in the gnotobiotic mouse experiment section. Two-way ANOVA was used to compare the means between groups.

### RNA-Seq

For biofilms in liquid culture, strains were grown as described previously for biofilm imaging. After 24 hours, the planktonic fraction of the wild-type strain was removed and cells were pelleted by centrifugation, whereas for the *ΔepsΔmfa* mutant, 2 ml of culture were directly pelleted. Cell pellets were immediately frozen in an ethanol-dry ice water and stored at -80°C. Samples for RNA-seq analysis from mice were obtained as described in the Gnotobiotic mouse section. RNA-Seq was performed at the DFI-MMF. Total RNA was extracted from three biological replicates per condition using the RNeasy Power Microbiome kit (Qiagen). Ribosomal RNA was depleted using the NEBNext rRNA Depletion Kit for bacteria (New England Biolabs), cDNA libraries were constructed using the NEB Ultra Directional RNA library prep kit for Illumina and sequenced on Illumina’s NextSeq 1000 platform at 2 × 100 bp read length. Transcriptomic profiles were analyzed as described previously^58^. Briefly, after quality trimming, reads were mapped to the PhvCL11 genome using Bowtie 2 (v. 2.4.2)^109–111^ and evaluated for differential gene expression using DESeq2 (v. 1.30.0)^112^ and edgeR (v. 3.32.1)^113^. If both statistical packages agreed (p_adj_ ≤ 0.05), a gene was considered differentially expressed.

### Plasmid transfer experiments

For plasmid transfer assays, an erythromycin resistance cassette was inserted in the intergenic region between INE91_04292 and INE91_04293 of pMMCAT in PhvCL11. A tetracycline resistance cassette was inserted in the genome of the recipient strain *B. uniformis* CL02T12C37 in the intergenic region between INE75_02569 and INE75_02570. Converse experiments were carried out where the donor was labeled with tetracycline and the recipient with erythromycin. For plasmid transfer assays on solid medium, strains were grown separately from an overnight culture, diluted 1:100 and incubated to an OD_600_ of 0.1. Next, 5 µL of each strain was mixed and spotted together on a BHIS plate. After 24 hours incubation, cells were retrieved from the plate and resuspended in 500 µL of basal medium for serial dilutions and plating in BHIS with erythromycin alone, with tetracycline alone, and with both antibiotics. To verify that the colonies arising on the double antibiotic plates were transconjugants, 10 colonies per plate were screened by PCR to verify that they were *B. uniformis* carrying pMMCAT (Table S7A). For conjugation experiments in biofilms, biofilms were grown as described in the mixed-species biofilm section, with both strains added at equal ratio. After mixing, plates were incubated anaerobically for 24 hours and biofilms were homogenized for CFU counting of donor, recipient and transformants on selective plates as before. Transfer experiments in mice were carried out as described in the Gnotobiotic Mouse section.

### Analysis of pMMCAT prevalence in sequenced genomes

We built a blastn database including all genomes from NCBI classified as belonging to the phylum Bacteroidota as of November 2023. We excluded genomes marked as suppressed or anomalous, Metagenomic Assembly Genomes (MAGs), and those that were identified as redundant or duplicated between Genbank and Refseq (Table S3). Metadata from each sample was parsed to follow a consistent format for country of isolation, year of isolation, host, and broad environmental categories for the isolation source. For large studies including multiple isolates from human cohorts, samples belonging to the same volunteer were identified from metadata and from supplementary tables in the relevant publications for each cohort. For the tallying of pMMCAT abundances, isolates of the same species cultured from the same volunteer were counted as a single entry, unless there were differences between them in pMMCAT blast results. We used the sequence of pMMCAT from *Bacteroides cellulosilyticus* CL06T03C01 (CP072253 plasmid pMMCAT_BcelCL06T03) as the query in these analyses. Genomes were considered to have pMMCAT if the hits covered more than 94% of the query sequence. A histogram plot of coverage in genomes with less than 94% coverage indicated a strong peak around 75% coverage. We downloaded the corresponding contigs and aligned them to the query in Snapgene (Dotmatics) using the Align to Reference tool to determine the degree of similarity.

### Multiple sequence alignment and phylogenetic reconstruction of pMMCAT sequences

For each strain that had pMMCAT, all contigs carrying high scoring segment pairs were downloaded. For heavily fragmented genomes, contigs were assembled using Ragout v2.3^114,115^ and the blast query sequence as reference. Genomes where some hits were part of very large contigs (e.g. chromosomal regions) or where scaffolding yielded patterns inconsistent with pMMCAT architecture were manually inspected. Quality of alignments to the reference (of both contigs and scaffolds) was visually checked on Snapgene using the Align to Reference tool. We determined that all cases where chromosomal hits were detected corresponded to lower identity matches of small pMMCAT regions (whereas non-chromosomal hits to pMMCAT are always >93% identical to the query). In the cases where scaffolding was inconsistent, all cases corresponded to integration of insertion sequence elements (ISE), other small mobile genetic elements or prophages in pMMCAT. These were spliced from the sequence for downstream analysis (the majority did not disrupt pMMCAT loci and were in intergenic regions). The 12 cases where integrated MGEs or ISEs interrupted important pMMCAT loci were excluded from further processing. Genome assemblies that were too fragmented or had too many gaps were excluded from the analysis. One of the genomes we sequenced from the Boston collection (*B. thetaiotaomicron* TS60) fell under this category but was kept in the analysis due to its historical importance, however the frequent gaps caused RaxML to artificially present this sequence as divergent.

All pMMCAT sequences (also including those from our newly sequenced genomes) were aligned to the reference in Snapgene to set to the same starting nucleotide, then the re-zeroed sequences were aligned in MAFFT v7.520^116^ with default parameters and the flag “--maxiterate 20 –reorder”. For strains from the same species isolated from the same person where the pMMCAT sequence was identical, only one sequence was kept for the following steps. Aligned sequences were visualized in the Integrative Genomics Viewer (IGV) 2.16.2^117^ to generate the coverage plots used to determine consensus and variability. A maximum likelihood phylogeny was generated in RaxML v 8.2.13^118^ using rapid bootstrapping, the GTRGAMMA model and autoMRE bootstrap convergence criterion. The resulting best tree was visualized in FigTree v1.4.4^119^. For analysis of *mfa* locus variation, alignments were inspected in Jalview 2.11.3.2^120^ and predicted Alphafold 3D structures^121^ of the base and stalk proteins were visualized in ChimeraX 1.7^122^.

### PCR screening of pMMCAT in clinical isolate collection and whole genome sequencing

Two collections of clinical Bacteroidales isolates from Boston area hospitals were provided by the laboratories of Lynn Bry and Andrew Onderdonk. Excluding *B. fragilis*, one collection included 18 isolates dating from 1980-1988 and the second one included 46 isolates from ∼1993-1997. Each strain was streaked for isolation and PCR was performed with each strain using Taq DNA polymerase (NEB) and the primers and reaction conditions listed in Table S7A to amplify one 610-bp region of the *eps* locus (NE91_04885), two regions of the *mfa* locus (540 bp and 505 bp), with the first region being very conserved in all *mfa* variants, and the complete 1483-bp region of the 16S RNA^19^. 16S RNA PCR products were sequenced via Sanger reaction at the University of Chicago DNA Sequencing Facility for species identification. For strains yielding *eps* or *mfa* PCR products, whole genome sequencing using hybrid Oxford Nanopore and Illumina was performed at the Duchossois Family Institute Microbiome Metagenomics facility (DFI-MMF, University of Chicago, Chicago, IL). For Illumina sequencing, bacterial samples were subjected to mechanical disruption on a bead mill homogenizer (Fisherbrand). DNA was purified using a spin column filter membrane and quantified using Qubit (Life Technologies). Libraries were prepared using 200 ng of genomic DNA and the QIAseq FX DNA library kit (Qiagen). The DNA was fragmented enzymatically, end repaired using T4 DNA polymerase, and Illumina compatible Unique Dual Index (UDI) adapters were added to the inserts. The prepared library was PCR amplified, recovered using magnetic beads, and QC was performed using Tapestation 4200 (Agilent Technologies). Libraries were sequenced on an NextSeq 1000 platform to generate 2 × 250 bp paired-end reads, producing approximately 5 million paired-end reads per sample. For Nanopore sequencing, DNA was extracted using the Monarch Genomic DNA Purification Kit (NEB), and quality checked using a genomic Tapestation 4200 (Agilent Technologies). Nanopore libraries were prepared using the Ligation Sequencing Kit (SQK-LSK109), the Native Barcoding Expansions 1-12 (EXP-NBD104) and 13-24 (EXP-NBD114), and the NEBNext Companion Module for Oxford Nanopore Technologies (E7180S). The shearing steps and first ethanol wash were eliminated to ensure high concentrations of long fragments. Using R9.4.1 flow cells, libraries were run on a MinION for 72 hours at ≈180 mV. Hybrid assemblies were completed using Unicycler (v0.4.8)^123^ with default parameters, using the non-filtered Nanopore long reads with and without prior trimming of the Illumina short reads as input. Where applicable, the short reads were trimmed by Trim Galore (v.0.4.5) with following parameters: the adapter and the leading 6 bp of forward and reverse short reads were trimmed off, the quality was controlled at 30, while minimum length was at 75 bp. Bandage v0.8.1^124^ was used to determine if reads mapped to a circular pMMCAT plasmid and estimate copy number. Prodigal 2.6.3^125^ was used for gene calls and Prokka 1.14.6^126^ was used for gene annotation.

### Metagenomic read recruitment, analysis of pMMCAT detection and quantification of promoter orientations

We analyzed 4,824 healthy human adult gut metagenomes and 215 children, listed on Table S7 along with their NCBI accession numbers and relevant metadata. These metagenomic studies comprised isolates from Australia (PRJEB6092), Austria^127^, Bangladesh^128^, Brazil^129^, Canada^130^, China^131,132^, Denmark^133^, England^134^, Ethiopia^76^, Fiji^15^, Finland^135^, Ghana^68^, India^136^, Israel^137^, Italy^77,138^, Japan^79^, Korea^139^, Madagascar^76^, Mongolia^76,140^, Netherlands^141^, Peru^142^, Spain^143^, Sweden^144,145^, Tanzania^68,77^, Thailand^2^ and the USA^2,83,142,146,147^. We also analyzed 1,283 gut metagenomes from infant-mother pairs from Italy^138^, Finland^135^, Sweden^144^ and the USA^146,148^, listed on Table S1F. 1430 metagenomes from IBD cohorts (Table S1E) from the USA^81–83^, Netherlands^81^ and Spain^80^ were analyzed. Other human fecal metagenomic studies included in the analysis were: two FMT cohorts of 32 patients with Chron’s disease^53^ or recurrent *C. difficile*^49^ (Table S1C); a challenge study of six volunteers to enterotoxigenic *E. coli* followed by ciprofloxacin treatment^51,52^ (Table S1B); a cohort of 12 healthy volunteers in Denmark who were treated with a 4-day course of combination antibiotics^50^ (Table S1A). 2972 human non-gut metagenomes (oral, nasopharyngeal, lung, skin, vaginal) available on the SRA were analyzed (Table S1H), as well as 4877 fecal samples from domesticated and wild animals (Table S1I, including pets, all common species of livestock, chimpanzees, gorillas and lab mice) and 10044 environmental samples (marine and freshwater environments, plants, soil, household and hospital surfaces). Metagenomic sewage samples analyzed were part of a global sewage survey^78^ that and included 950 samples from 101 countries and 242 cities (Table S1F). Finally, archeological metagenomic projects analyzed included two samples of medieval coprolites/latrine sediments (Latvia and Israel)^66^, the colon contents of the Copper Age iceman (Ötzi)^67,68^, pre-Columbian coprolites in America (12 samples from 500-600 A.D., Mexico^69,70^ and Puerto Rico^71^) and Neanderthal coprolites (7 samples from Spain)^72^.

Metagenomic analyses were performed using the anvi’o v7 software suite^149^. Raw reads were quality-filtered using illumina-utils v1.4.4 program ’iu-filter-quality-minochè^150^ with default parameters. Bowtie2 v2.5.1^109–111^ was used to map reads to the pMMCAT sequence from *Bacteroides cellulosilyticus* CL06T03C01 or the different promoter orientation templates, using parameters “--end-to-end -D 25 -R 5 -N 1 -L 20 -i S,1,0.50” for pMMCAT and “--end-to-end –fast” for the promoter orientation templates. Samtools v1.18^151^ was used to convert resulting SAM files into sorted and indexed BAM files. Anvi’o contigs databases for each mapping template were generated using the command ’anvi-gen-contigs-databasè. For each mapped metagenome, anvi’o profile databases (coverage, detection and SNV statistics) were generated using ’anvi-profile --cluster’. For each dataset, profiles from all samples were combined using ’anvi-mergè. Coverage and detection statistics for each merged profile database, were recovered using ’anvi-summarize --init-gene-coverages’ flag. Samples were counted as having pMMCAT if they had a detection value larger than 0.94. Kraken 2.0.8-beta^152^ was used to estimate the taxonomic composition of each metagenome.

## REFERENCES

1. Wexler, A.G., and Goodman, A.L. (2017). An insider’s perspective: Bacteroides as a window into the microbiome. Nature Microbiology 2, 17026. 10.1038/nmicrobiol.2017.26.

2. Vangay, P., Johnson, A.J., Ward, T.L., Al-Ghalith, G.A., Shields-Cutler, R.R., Hillmann, B.M., Lucas, S.K., Beura, L.K., Thompson, E.A., Till, L.M., et al. (2018). US Immigration Westernizes the Human Gut Microbiome. Cell 175, 962–972.e10. 10.1016/j.cell.2018.10.029.

3. Faith, J.J., Guruge, J.L., Charbonneau, M., Subramanian, S., Seedorf, H., Goodman, A.L., Clemente, J.C., Knight, R., Heath, A.C., Leibel, R.L., et al. (2013). The Long-Term Stability of the Human Gut Microbiota. Science 341, 1237439–1237439. 10.1126/science.1237439.

4. Mehta, R.S., Abu-Ali, G.S., Drew, D.A., Lloyd-Price, J., Subramanian, A., Lochhead, P., Joshi, A.D., Ivey, K.L., Khalili, H., Brown, G.T., et al. (2018). Stability of the human faecal microbiome in a cohort of adult men. Nature Microbiology 3, 347–355. 10.1038/s41564-017-0096-0.

5. Moya, A., and Ferrer, M. (2016). Functional Redundancy-Induced Stability of Gut Microbiota Subjected to Disturbance. Trends in Microbiology 24, 402–413. 10.1016/j.tim.2016.02.002.

6. Lozupone, C.A., Stombaugh, J.I., Gordon, J.I., Jansson, J.K., and Knight, R. (2012). Diversity, stability and resilience of the human gut microbiota. Nature 489, 220–230. 10.1038/nature11550.

7. Sommer, F., Anderson, J.M., Bharti, R., Raes, J., and Rosenstiel, P. (2017). The resilience of the intestinal microbiota influences health and disease. Nature Reviews Microbiology 15, 630–638. 10.1038/nrmicro.2017.58.

8. Lokmer, A., Aflalo, S., Amougou, N., Lafosse, S., Froment, A., Tabe, F.E., Poyet, M., Groussin, M., Said-Mohamed, R., and Ségurel, L. (2020). Response of the human gut and saliva microbiome to urbanization in Cameroon. Sci Rep 10, 2856. 10.1038/s41598-020-59849-9.

9. Coyne, M.J., Zitomersky, N.L., McGuire, A.M., Earl, A.M., and Comstock, L.E. (2014). Evidence of Extensive DNA Transfer between Bacteroidales Species within the Human Gut. mBio 5, e01305-14–e01305-14. 10.1128/mBio.01305-14.

10. Groussin, M., Poyet, M., Sistiaga, A., Kearney, S.M., Moniz, K., Noel, M., Hooker, J., Gibbons, S.M., Segurel, L., Froment, A., et al. (2021). Elevated rates of horizontal gene transfer in the industrialized human microbiome. Cell 184, 2053–2067.e18. 10.1016/j.cell.2021.02.052.

11. Jiang, X., Hall, A.B., Xavier, R.J., and Alm, E.J. (2019). Comprehensive analysis of chromosomal mobile genetic elements in the gut microbiome reveals phylum-level niche-adaptive gene pools. PLOS ONE 14, e0223680. 10.1371/journal.pone.0223680.

12. Kent, A.G., Vill, A.C., Shi, Q., Satlin, M.J., and Brito, I.L. (2020). Widespread transfer of mobile antibiotic resistance genes within individual gut microbiomes revealed through bacterial Hi-C. Nat Commun 11, 4379. 10.1038/s41467-020-18164-7.

13. Yaffe, E., and Relman, D.A. (2020). Tracking microbial evolution in the human gut using Hi-C reveals extensive horizontal gene transfer, persistence and adaptation. Nature Microbiology 5, 343–353. 10.1038/s41564-019-0625-0.

14. Zhou, H., Beltrán, J.F., and Brito, I.L. (2021). Functions predict horizontal gene transfer and the emergence of antibiotic resistance. Sci. Adv. 7, eabj5056. 10.1126/sciadv.abj5056.

15. Brito, I.L., Yilmaz, S., Huang, K., Xu, L., Jupiter, S.D., Jenkins, A.P., Naisilisili, W., Tamminen, M., Smillie, C.S., Wortman, J.R., et al. (2016). Mobile genes in the human microbiome are structured from global to individual scales. Nature 535, 435–439. 10.1038/nature18927.

16. García-Bayona, L., Coyne, M.J., and Comstock, L.E. (2021). Mobile Type VI secretion system loci of the gut Bacteroidales display extensive intra-ecosystem transfer, multi-species spread and geographical clustering. PLoS Genet 17, e1009541. 10.1371/journal.pgen.1009541.

17. Conteville, L.C., and Vicente, A.C.P. (2022). A plasmid network from the gut microbiome of semi-isolated human groups reveals unique and shared metabolic and virulence traits. Sci Rep 12, 12102. 10.1038/s41598-022-16392-z.

18. Poyet, M., Groussin, M., Gibbons, S.M., Avila-Pacheco, J., Jiang, X., Kearney, S.M., Perrotta, A.R., Berdy, B., Zhao, S., Lieberman, T.D., et al. (2019). A library of human gut bacterial isolates paired with longitudinal multiomics data enables mechanistic microbiome research. Nat Med 25, 1442– 1452. 10.1038/s41591-019-0559-3.

19. Zitomersky, N.L., Coyne, M.J., and Comstock, L.E. (2011). Longitudinal Analysis of the Prevalence, Maintenance, and IgA Response to Species of the Order Bacteroidales in the Human Gut. Infection and Immunity 79, 2012–2020. 10.1128/IAI.01348-10.

20. Browne, H.P., Forster, S.C., Anonye, B.O., Kumar, N., Neville, B.A., Stares, M.D., Goulding, D., and Lawley, T.D. (2016). Culturing of “unculturable” human microbiota reveals novel taxa and extensive sporulation. Nature 533, 543–546. 10.1038/nature17645.

21. Zou, Y., Xue, W., Luo, G., Deng, Z., Qin, P., Guo, R., Sun, H., Xia, Y., Liang, S., Dai, Y., et al. (2019). 1,520 reference genomes from cultivated human gut bacteria enable functional microbiome analyses. Nat Biotechnol 37, 179–185. 10.1038/s41587-018-0008-8.

22. Peed, L., Parker, A.C., and Smith, C.J. (2010). Genetic and Functional Analyses of the mob Operon on Conjugative Transposon CTn341 from Bacteroides spp. Journal of Bacteriology 192, 4643–4650. 10.1128/JB.00317-10.

23. Bacic, M., Parker, A.C., Stagg, J., Whitley, H.P., Wells, W.G., Jacob, L.A., and Smith, C.J. (2005). Genetic and Structural Analysis of the Bacteroides Conjugative Transposon CTn341. Journal of Bacteriology 187, 2858–2869. 10.1128/JB.187.8.2858-2869.2005.

24. Waters, J.L., Wang, G.-R., and Salyers, A.A. (2013). Tetracycline-Related Transcriptional Regulation of the CTnDOT Mobilization Region. Journal of Bacteriology 195, 5431–5438. 10.1128/JB.00691-13.

25. Coyne, M.J., Roelofs, K.G., and Comstock, L.E. (2016). Type VI secretion systems of human gut Bacteroidales segregate into three genetic architectures, two of which are contained on mobile genetic elements. BMC Genomics 17. 10.1186/s12864-016-2377-z.

26. Burmølle, M., Ren, D., Bjarnsholt, T., and Sørensen, S.J. (2014). Interactions in multispecies biofilms: do they actually matter? Trends in Microbiology 22, 84–91. 10.1016/j.tim.2013.12.004.

27. Elias, S., and Banin, E. (2012). Multi-species biofilms: living with friendly neighbors. FEMS Microbiology Reviews 36, 990–1004. 10.1111/j.1574-6976.2012.00325.x.

28. Tomkovich, S., Dejea, C.M., Winglee, K., Drewes, J.L., Chung, L., Housseau, F., Pope, J.L., Gauthier, J., Sun, X., Mühlbauer, M., et al. (2019). Human colon mucosal biofilms from healthy or colon cancer hosts are carcinogenic. Journal of Clinical Investigation 129, 1699–1712. 10.1172/JCI124196.

29. Domingue, J.C., Drewes, J.L., Merlo, C.A., Housseau, F., and Sears, C.L. (2020). Host responses to mucosal biofilms in the lung and gut. Mucosal Immunology 13, 413–422. 10.1038/s41385-020-0270-1.

30. Béchon, N., and Ghigo, J.-M. (2022). Gut biofilms: **Bacteroides** as model symbionts to study biofilm formation by intestinal anaerobes. FEMS Microbiology Reviews 46, fuab054. 10.1093/femsre/fuab054.

31. Wong, J.P.H., Fischer-Stettler, M., Zeeman, S.C., Battin, T.J., and Persat, A. (2023). Fluid flow structures gut microbiota biofilm communities by distributing public goods. Proc. Natl. Acad. Sci. U.S.A. 120, e2217577120. 10.1073/pnas.2217577120.

32. Dragoš, A., and Kovács, Á.T. (2017). The Peculiar Functions of the Bacterial Extracellular Matrix. Trends Microbiol 25, 257–266. 10.1016/j.tim.2016.12.010.

33. Oliveira, R.A., Cabral, V., Torcato, I., and Xavier, K.B. (2023). Deciphering the quorum-sensing lexicon of the gut microbiota. Cell Host Microbe 31, 500–512. 10.1016/j.chom.2023.03.015.

34. Jin, X., and Marshall, J.S. (2020). Mechanics of biofilms formed of bacteria with fimbriae appendages. PLoS One 15, e0243280. 10.1371/journal.pone.0243280.

35. Baier, F., and Tokuriki, N. (2014). Connectivity between Catalytic Landscapes of the Metallo-β-Lactamase Superfamily. Journal of Molecular Biology 426, 2442–2456. 10.1016/j.jmb.2014.04.013.

36. Latham, J.A., Barr, I., and Klinman, J.P. (2017). At the confluence of ribosomally synthesized peptide modification and radical **S** -adenosylmethionine (SAM) enzymology. Journal of Biological Chemistry 292, 16397–16405. 10.1074/jbc.R117.797399.

37. Béchon, N., Mihajlovic, J., Lopes, A.-A., Vendrell-Fernández, S., Deschamps, J., Briandet, R., Sismeiro, O., Martin-Verstraete, I., Dupuy, B., and Ghigo, J.-M. (2022). *Bacteroides thetaiotaomicron* uses a widespread extracellular DNase to promote bile-dependent biofilm formation. Proc. Natl. Acad. Sci. U.S.A. 119, e2111228119. 10.1073/pnas.2111228119.

38. Mihajlovic, J., Bechon, N., Ivanova, C., Chain, F., Almeida, A., Langella, P., Beloin, C., and Ghigo, J.-M. (2019). A Putative Type V Pilus Contributes to *Bacteroides thetaiotaomicron* Biofilm Formation Capacity. Journal of Bacteriology 201. 10.1128/JB.00650-18.

39. Krinos, C.M., Coyne, M.J., Weinacht, K.G., Tzianabos, A.O., Kasper, D.L., and Comstock, L.E. (2001). Extensive surface diversity of a commensal microorganism by multiple DNA inversions. Nature 414, 555–558. 10.1038/35107092.

40. Lan, F., Saba, J., Ross, T.D., Zhou, Z., Krauska, K., Anantharaman, K., Landick, R., and Venturelli, O.S. (2024). Massively parallel single-cell sequencing of diverse microbial populations. Nat Methods. 10.1038/s41592-023-02157-7.

41. Porter, N.T., Hryckowian, A.J., Merrill, B.D., Fuentes, J.J., Gardner, J.O., Glowacki, R.W.P., Singh, S., Crawford, R.D., Snitkin, E.S., Sonnenburg, J.L., et al. (2020). Phase-variable capsular polysaccharides and lipoproteins modify bacteriophage susceptibility in Bacteroides thetaiotaomicron. Nat Microbiol 5, 1170–1181. 10.1038/s41564-020-0746-5.

42. Fletcher, C.M., Coyne, M.J., Bentley, D.L., Villa, O.F., and Comstock, L.E. (2007). Phase-variable expression of a family of glycoproteins imparts a dynamic surface to a symbiont in its human intestinal ecosystem. Proc. Natl. Acad. Sci. U.S.A. 104, 2413–2418. 10.1073/pnas.0608797104.

43. DePas, W.H., Starwalt-Lee, R., Van Sambeek, L., Ravindra Kumar, S., Gradinaru, V., and Newman, D.K. (2016). Exposing the Three-Dimensional Biogeography and Metabolic States of Pathogens in Cystic Fibrosis Sputum via Hydrogel Embedding, Clearing, and rRNA Labeling. mBio 7. 10.1128/mBio.00796-16.

44. Gallego-Hernandez, A.L., DePas, W.H., Park, J.H., Teschler, J.K., Hartmann, R., Jeckel, H., Drescher, K., Beyhan, S., Newman, D.K., and Yildiz, F.H. (2020). Upregulation of virulence genes promotes *Vibrio cholerae* biofilm hyperinfectivity. Proceedings of the National Academy of Sciences 117, 11010–11017. 10.1073/pnas.1916571117.

45. Abed, J., Emgård, J.E.M., Zamir, G., Faroja, M., Almogy, G., Grenov, A., Sol, A., Naor, R., Pikarsky, E., Atlan, K.A., et al. (2016). Fap2 Mediates Fusobacterium nucleatum Colorectal Adenocarcinoma Enrichment by Binding to Tumor-Expressed Gal-GalNAc. Cell Host & Microbe 20, 215–225. 10.1016/j.chom.2016.07.006.

46. White, M.T., and Sears, C.L. (2023). The microbial landscape of colorectal cancer. Nat Rev Microbiol. 10.1038/s41579-023-00973-4.

47. Coyne, M.J., Béchon, N., Matano, L.M., McEneany, V.L., Chatzidaki-Livanis, M., and Comstock, L.E. (2019). A family of anti-Bacteroidales peptide toxins wide-spread in the human gut microbiota. Nature Communications 10. 10.1038/s41467-019-11494-1.

48. Fagarasan, S., Muramatsu, M., Suzuki, K., Nagaoka, H., Hiai, H., and Honjo, T. (2002). Critical Roles of Activation-Induced Cytidine Deaminase in the Homeostasis of Gut Flora. Science 298, 1424– 1427. 10.1126/science.1077336.

49. Smillie, C.S., Sauk, J., Gevers, D., Friedman, J., Sung, J., Youngster, I., Hohmann, E.L., Staley, C., Khoruts, A., Sadowsky, M.J., et al. (2018). Strain Tracking Reveals the Determinants of Bacterial Engraftment in the Human Gut Following Fecal Microbiota Transplantation. Cell Host & Microbe 23, 229–240.e5. 10.1016/j.chom.2018.01.003.

50. Palleja, A., Mikkelsen, K.H., Forslund, S.K., Kashani, A., Allin, K.H., Nielsen, T., Hansen, T.H., Liang, S., Feng, Q., Zhang, C., et al. (2018). Recovery of gut microbiota of healthy adults following antibiotic exposure. Nat Microbiol 3, 1255–1265. 10.1038/s41564-018-0257-9.

51. Tvedte, E.S., Michalski, J., Cheng, S., Patkus, R.S., Tallon, L.J., Sadzewicz, L., Bruno, V.M., Silva, J.C., Rasko, D.A., and Dunning Hotopp, J.C. (2021). Evaluation of a high-throughput, cost-effective Illumina library preparation kit. Sci Rep 11, 15925. 10.1038/s41598-021-94911-0.

52. McArthur, M.A., Chen, W.H., Magder, L., Levine, M.M., and Sztein, M.B. (2017). Impact of CD4+ T Cell Responses on Clinical Outcome following Oral Administration of Wild-Type Enterotoxigenic Escherichia coli in Humans. PLoS Negl Trop Dis 11, e0005291. 10.1371/journal.pntd.0005291.

53. Vaughn, B.P., Vatanen, T., Allegretti, J.R., Bai, A., Xavier, R.J., Korzenik, J., Gevers, D., Ting, A., Robson, S.C., and Moss, A.C. (2016). Increased Intestinal Microbial Diversity Following Fecal Microbiota Transplant for Active Crohn’s Disease. Inflamm Bowel Dis 22, 2182–2190. 10.1097/MIB.0000000000000893.

54. Alauzet, C., Lozniewski, A., and Marchandin, H. (2019). Metronidazole resistance and nim genes in anaerobes: A review. Anaerobe 55, 40–53. 10.1016/j.anaerobe.2018.10.004.

55. Humphries, J., Xiong, L., Liu, J., Prindle, A., Yuan, F., Arjes, H.A., Tsimring, L., and Süel, G.M. (2017). Species-Independent Attraction to Biofilms through Electrical Signaling. Cell 168, 200–209.e12. 10.1016/j.cell.2016.12.014.

56. Van Espen, L., Bak, E.G., Beller, L., Close, L., Deboutte, W., Juel, H.B., Nielsen, T., Sinar, D., De Coninck, L., Frithioff-Bøjsøe, C., et al. (2021). A Previously Undescribed Highly Prevalent Phage Identified in a Danish Enteric Virome Catalog. mSystems 6, e00382–21. 10.1128/mSystems.00382-21.

57. Fogarty, E.C., Schechter, M.S., Lolans, K., Sheahan, M.L., Veseli, I., Moore, R.M., Kiefl, E., Moody, T., Rice, P.A., Yu, M.K., et al. (2024). A cryptic plasmid is among the most numerous genetic elements in the human gut. Cell 187, 1206–1222.e16. 10.1016/j.cell.2024.01.039.

58. Sheahan, M.L., Coyne, M.J., Flores, K., Garcia-Bayona, L., Chatzidaki-Livanis, M., Sundararajan, A., Holst, A.Q., Barquera, B., and Comstock, L.E. (2023). A ubiquitous mobile genetic element disarms a bacterial antagonist of the gut microbiota (Microbiology) 10.1101/2023.08.25.553775.

59. Stevens, A.M., Shoemaker, N.B., and Salyers, A.A. (1990). The region of a Bacteroides conjugal chromosomal tetracycline resistance element which is responsible for production of plasmidlike forms from unlinked chromosomal DNA might also be involved in transfer of the element. J Bacteriol 172, 4271–4279. 10.1128/jb.172.8.4271-4279.1990.

60. Blanco-Míguez, A., Gálvez, E.J.C., Pasolli, E., De Filippis, F., Amend, L., Huang, K.D., Manghi, P., Lesker, T.-R., Riedel, T., Cova, L., et al. (2023). Extension of the Segatella copri complex to 13 species with distinct large extrachromosomal elements and associations with host conditions. Cell Host Microbe 31, 1804–1819.e9. 10.1016/j.chom.2023.09.013.

61. Mitchell, S., Bull, M., Muscatello, G., Chapman, B., and Coleman, N.V. (2021). The equine hindgut as a reservoir of mobile genetic elements and antimicrobial resistance genes. Critical Reviews in Microbiology 47, 543–561. 10.1080/1040841X.2021.1907301.

62. Wang, C., Song, Y., Tang, N., Zhang, G., Leclercq, S.O., and Feng, J. (2021). The Shared Resistome of Human and Pig Microbiota Is Mobilized by Distinct Genetic Elements. Appl Environ Microbiol 87, e01910–20. 10.1128/AEM.01910-20.

63. Ma, T., McAllister, T.A., and Guan, L.L. (2021). A review of the resistome within the digestive tract of livestock. J Animal Sci Biotechnol 12, 121. 10.1186/s40104-021-00643-6.

64. Delaney, M.L., and Onderdonk, A.B. (1997). Evaluation of the AnaeroPack system for growth of clinically significant anaerobes. J Clin Microbiol 35, 558–562. 10.1128/jcm.35.3.558-562.1997.

65. Tisza, M.J., Smith, D.D.N., Clark, A.E., Youn, J.-H., NISC Comparative Sequencing Program, Khil, P.P., and Dekker, J.P. (2023). Roving methyltransferases generate a mosaic epigenetic landscape and influence evolution in Bacteroides fragilis group. Nat Commun 14, 4082. 10.1038/s41467-023-39892-6.

66. Sabin, S., Yeh, H.-Y., Pluskowski, A., Clamer, C., Mitchell, P.D., and Bos, K.I. (2020). Estimating molecular preservation of the intestinal microbiome via metagenomic analyses of latrine sediments from two medieval cities. Phil. Trans. R. Soc. B 375, 20190576. 10.1098/rstb.2019.0576.

67. Lugli, G.A., Milani, C., Mancabelli, L., Turroni, F., Ferrario, C., Duranti, S., Van Sinderen, D., and Ventura, M. (2017). Ancient bacteria of the Ötzi’s microbiome: a genomic tale from the Copper Age. Microbiome 5, 5. 10.1186/s40168-016-0221-y.

68. Tett, A., Huang, K.D., Asnicar, F., Fehlner-Peach, H., Pasolli, E., Karcher, N., Armanini, F., Manghi, P., Bonham, K., Zolfo, M., et al. (2019). The Prevotella copri Complex Comprises Four Distinct Clades Underrepresented in Westernized Populations. Cell Host & Microbe 26, 666–679.e7. 10.1016/j.chom.2019.08.018.

69. Hagan, R.W., Hofman, C.A., Hübner, A., Reinhard, K., Schnorr, S., Lewis, C.M., Sankaranarayanan, K., and Warinner, C.G. (2020). Comparison of extraction methods for recovering ancient microbial DNA from paleofeces. American J Phys Anthropol 171, 275–284. 10.1002/ajpa.23978.

70. Borry, M., Cordova, B., Perri, A., Wibowo, M., Prasad Honap, T., Ko, J., Yu, J., Britton, K., Girdland-Flink, L., Power, R.C., et al. (2020). CoproID predicts the source of coprolites and paleofeces using microbiome composition and host DNA content. PeerJ 8, e9001. 10.7717/peerj.9001.

71. Reynoso-García, J., Santiago-Rodriguez, T.M., Narganes-Storde, Y., Cano, R.J., and Toranzos, G.A. (2023). Edible flora in pre-Columbian Caribbean coprolites: Expected and unexpected data. PLoS ONE 18, e0292077. 10.1371/journal.pone.0292077.

72. Rampelli, S., Turroni, S., Mallol, C., Hernandez, C., Galván, B., Sistiaga, A., Biagi, E., Astolfi, A., Brigidi, P., Benazzi, S., et al. (2021). Components of a Neanderthal gut microbiome recovered from fecal sediments from El Salt. Commun Biol 4, 169. 10.1038/s42003-021-01689-y.

73. Hasegawa, Y., and Nagano, K. (2021). Porphyromonas gingivalis FimA and Mfa1 fimbriae: Current insights on localization, function, biogenesis, and genotype. Japanese Dental Science Review 57, 190–200. 10.1016/j.jdsr.2021.09.003.

74. Aggarwala, V., Mogno, I., Li, Z., Yang, C., Britton, G.J., Chen-Liaw, A., Mitcham, J., Bongers, G., Gevers, D., Clemente, J.C., et al. (2021). Precise quantification of bacterial strains after fecal microbiota transplantation delineates long-term engraftment and explains outcomes. Nat Microbiol 6, 1309–1318. 10.1038/s41564-021-00966-0.

75. Zhang, Z.J., Cole, C.G., Coyne, M.J., Lin, H., Dylla, N., Smith, R.C., Waligurski, E., Ramaswamy, R., Woodson, C., Burgo, V., et al. (2024). Comprehensive analyses of a large human gut Bacteroidales culture collection reveal species and strain level diversity and evolution. Preprint, 10.1101/2024.03.08.584156 10.1101/2024.03.08.584156.

76. Pasolli, E., Asnicar, F., Manara, S., Zolfo, M., Karcher, N., Armanini, F., Beghini, F., Manghi, P., Tett, A., Ghensi, P., et al. (2019). Extensive Unexplored Human Microbiome Diversity Revealed by Over 150,000 Genomes from Metagenomes Spanning Age, Geography, and Lifestyle. Cell 176, 649–662.e20. 10.1016/j.cell.2019.01.001.

77. Rampelli, S., Schnorr, S.L., Consolandi, C., Turroni, S., Severgnini, M., Peano, C., Brigidi, P., Crittenden, A.N., Henry, A.G., and Candela, M. (2015). Metagenome Sequencing of the Hadza Hunter-Gatherer Gut Microbiota. Curr Biol 25, 1682–1693. 10.1016/j.cub.2015.04.055.

78. Munk, P., Brinch, C., Møller, F.D., Petersen, T.N., Hendriksen, R.S., Seyfarth, A.M., Kjeldgaard, J.S., Svendsen, C.A., Van Bunnik, B., Berglund, F., et al. (2022). Genomic analysis of sewage from 101 countries reveals global landscape of antimicrobial resistance. Nat Commun 13, 7251. 10.1038/s41467-022-34312-7.

79. Yachida, S., Mizutani, S., Shiroma, H., Shiba, S., Nakajima, T., Sakamoto, T., Watanabe, H., Masuda, K., Nishimoto, Y., Kubo, M., et al. (2019). Metagenomic and metabolomic analyses reveal distinct stage-specific phenotypes of the gut microbiota in colorectal cancer. Nat Med 25, 968–976. 10.1038/s41591-019-0458-7.

80. MetaHIT Consortium, Qin, J., Li, R., Raes, J., Arumugam, M., Burgdorf, K.S., Manichanh, C., Nielsen, T., Pons, N., Levenez, F., et al. (2010). A human gut microbial gene catalogue established by metagenomic sequencing. Nature 464, 59–65. 10.1038/nature08821.

81. Franzosa, E.A., Sirota-Madi, A., Avila-Pacheco, J., Fornelos, N., Haiser, H.J., Reinker, S., Vatanen, T., Hall, A.B., Mallick, H., McIver, L.J., et al. (2019). Gut microbiome structure and metabolic activity in inflammatory bowel disease. Nature Microbiology 4, 293–305. 10.1038/s41564-018-0306-4.

82. Schirmer, M., Franzosa, E.A., Lloyd-Price, J., McIver, L.J., Schwager, R., Poon, T.W., Ananthakrishnan, A.N., Andrews, E., Barron, G., Lake, K., et al. (2018). Dynamics of metatranscription in the inflammatory bowel disease gut microbiome. Nature Microbiology 3, 337–346. 10.1038/s41564-017-0089-z.

83. Lloyd-Price, J., Arze, C., Ananthakrishnan, A.N., Schirmer, M., Avila-Pacheco, J., Poon, T.W., Andrews, E., Ajami, N.J., Bonham, K.S., Brislawn, C.J., et al. (2019). Multi-omics of the gut microbial ecosystem in inflammatory bowel diseases. Nature 569, 655–662. 10.1038/s41586-019-1237-9.

84. Chng, K.R., Li, C., Bertrand, D., Ng, A.H.Q., Kwah, J.S., Low, H.M., Tong, C., Natrajan, M., Zhang, M.H., Xu, L., et al. (2020). Cartography of opportunistic pathogens and antibiotic resistance genes in a tertiary hospital environment. Nat Med 26, 941–951. 10.1038/s41591-020-0894-4.

85. Hall, R.J., Whelan, F.J., McInerney, J.O., Ou, Y., and Domingo-Sananes, M.R. (2020). Horizontal Gene Transfer as a Source of Conflict and Cooperation in Prokaryotes. Front. Microbiol. 11, 1569. 10.3389/fmicb.2020.01569.

86. Dimitriu, T., Lotton, C., Bénard-Capelle, J., Misevic, D., Brown, S.P., Lindner, A.B., and Taddei, F. (2014). Genetic information transfer promotes cooperation in bacteria. Proc. Natl. Acad. Sci. U.S.A. 111, 11103–11108. 10.1073/pnas.1406840111.

87. Mc Ginty, S.E., Rankin, D.J., and Brown, S.P. (2011). Horizontal Gene Transfer and the Evolution of Bacterial Cooperation: Mobile Elements and Bacterial Cooperation. Evolution 65, 21–32. 10.1111/j.1558-5646.2010.01121.x.

88. Dewar, A.E., Thomas, J.L., Scott, T.W., Wild, G., Griffin, A.S., West, S.A., and Ghoul, M. (2021). Plasmids do not consistently stabilize cooperation across bacteria but may promote broad pathogen host-range. Nat Ecol Evol 5, 1624–1636. 10.1038/s41559-021-01573-2.

89. Lee, I.P.A., Eldakar, O.T., Gogarten, J.P., and Andam, C.P. (2022). Bacterial cooperation through horizontal gene transfer. Trends in Ecology & Evolution 37, 223–232. 10.1016/j.tree.2021.11.006.

90. Coyte, K.Z., and Rakoff-Nahoum, S. (2019). Understanding Competition and Cooperation within the Mammalian Gut Microbiome. Curr Biol 29, R538–R544. 10.1016/j.cub.2019.04.017.

91. Simonet, C., and McNally, L. (2021). Kin selection explains the evolution of cooperation in the gut microbiota. Proc Natl Acad Sci U S A 118. 10.1073/pnas.2016046118.

92. Ghigo, J.-M. (2001). Natural conjugative plasmids induce bacterial biofilm development. Nature 412, 442–445. 10.1038/35086581.

93. Gama, J.A., Fredheim, E.G.A., Cléon, F., Reis, A.M., Zilhão, R., and Dionisio, F. (2020). Dominance Between Plasmids Determines the Extent of Biofilm Formation. Front. Microbiol. 11, 2070. 10.3389/fmicb.2020.02070.

94. Nakao, R., Myint, S.L., Wai, S.N., and Uhlin, B.E. (2018). Enhanced Biofilm Formation and Membrane Vesicle Release by Escherichia coli Expressing a Commonly Occurring Plasmid Gene, kil. Front. Microbiol. 9, 2605. 10.3389/fmicb.2018.02605.

95. Tytgat, H.L.P., Nobrega, F.L., van der Oost, J., and de Vos, W.M. (2019). Bowel Biofilms: Tipping Points between a Healthy and Compromised Gut? Trends in Microbiology 27, 17–25. 10.1016/j.tim.2018.08.009.

96. Béchon, N., Mihajlovic, J., Vendrell-Fernández, S., Chain, F., Langella, P., Beloin, C., and Ghigo, J.-M. (2020). Capsular Polysaccharide Cross-Regulation Modulates *Bacteroides thetaiotaomicron* Biofilm Formation. mBio 11. 10.1128/mBio.00729-20.

97. Weinacht, K.G., Roche, H., Krinos, C.M., Coyne, M.J., Parkhill, J., and Comstock, L.E. (2004). Tyrosine site-specific recombinases mediate DNA inversions affecting the expression of outer surface proteins of Bacteroides fragilis: DNA inversions regulate surface proteins of B. fragilis. Molecular Microbiology 53, 1319–1330. 10.1111/j.1365-2958.2004.04219.x.

98. Choi, H.M.T., Schwarzkopf, M., Fornace, M.E., Acharya, A., Artavanis, G., Stegmaier, J., Cunha, A., and Pierce, N.A. (2018). Third-generation *in situ* hybridization chain reaction: multiplexed, quantitative, sensitive, versatile, robust. Development 145, dev165753. 10.1242/dev.165753.

99. Røder, H.L., Olsen, N.M.C., Whiteley, M., and Burmølle, M. (2020). Unravelling interspecies interactions across heterogeneities in complex biofilm communities. Environmental Microbiology 22, 5–16. 10.1111/1462-2920.14834.

100. De Lorenzo, V., and Timmis, K.N. (1994). [31] Analysis and construction of stable phenotypes in gram-negative bacteria with Tn5- and Tn10-derived minitransposons. In Methods in Enzymology (Elsevier), pp. 386–405. 10.1016/0076-6879(94)35157-0.

101. García-Bayona, L., and Comstock, L.E. (2019). Streamlined Genetic Manipulation of Diverse *Bacteroides* and *Parabacteroides* Isolates from the Human Gut Microbiota. mBio 10. 10.1128/mBio.01762-19.

102. Chatzidaki-Livanis, M., Geva-Zatorsky, N., and Comstock, L.E. (2016). *Bacteroides fragilis* type VI secretion systems use novel effector and immunity proteins to antagonize human gut Bacteroidales species. Proceedings of the National Academy of Sciences 113, 3627–3632. 10.1073/pnas.1522510113.

103. Studier, F.W., and Moffatt, B.A. (1986). Use of bacteriophage T7 RNA polymerase to direct selective high-level expression of cloned genes. J Mol Biol 189, 113–130. 10.1016/0022-2836(86)90385-2.

104. Chatzidaki-Livanis, M., Coyne, M.J., Roche-Hakansson, H., and Comstock, L.E. (2008). Expression of a Uniquely Regulated Extracellular Polysaccharide Confers a Large-Capsule Phenotype to Bacteroides fragilis. Journal of Bacteriology 190, 1020–1026. 10.1128/JB.01519-07.

105. Kwasny, S.M., and Opperman, T.J. (2010). Static biofilm cultures of Gram-positive pathogens grown in a microtiter format used for anti-biofilm drug discovery. Curr Protoc Pharmacol Chapter 13, Unit 13A.8. 10.1002/0471141755.ph13a08s50.

106. Yang, B., Treweek, J.B., Kulkarni, R.P., Deverman, B.E., Chen, C.-K., Lubeck, E., Shah, S., Cai, L., and Gradinaru, V. (2014). Single-Cell Phenotyping within Transparent Intact Tissue through Whole-Body Clearing. Cell 158, 945–958. 10.1016/j.cell.2014.07.017.

107. Bolger, A.M., Lohse, M., and Usadel, B. (2014). Trimmomatic: a flexible trimmer for Illumina sequence data. Bioinformatics 30, 2114–2120. 10.1093/bioinformatics/btu170.

108. Blanco-Míguez, A., Beghini, F., Cumbo, F., McIver, L.J., Thompson, K.N., Zolfo, M., Manghi, P., Dubois, L., Huang, K.D., Thomas, A.M., et al. (2023). Extending and improving metagenomic taxonomic profiling with uncharacterized species using MetaPhlAn 4. Nat Biotechnol 41, 1633– 1644. 10.1038/s41587-023-01688-w.

109. Langmead, B., Wilks, C., Antonescu, V., and Charles, R. (2019). Scaling read aligners to hundreds of threads on general-purpose processors. Bioinformatics 35, 421–432. 10.1093/bioinformatics/bty648.

110. Langmead, B., and Salzberg, S.L. (2012). Fast gapped-read alignment with Bowtie 2. Nat Methods 9, 357–359. 10.1038/nmeth.1923.

111. Langmead, B., Trapnell, C., Pop, M., and Salzberg, S.L. (2009). Ultrafast and memory-efficient alignment of short DNA sequences to the human genome. Genome Biology 10, R25. 10.1186/gb-2009-10-3-r25.

112. Love, M.I., Huber, W., and Anders, S. (2014). Moderated estimation of fold change and dispersion for RNA-seq data with DESeq2. Genome Biol 15, 550. 10.1186/s13059-014-0550-8.

113. Robinson, M.D., McCarthy, D.J., and Smyth, G.K. (2010). edgeR: a Bioconductor package for differential expression analysis of digital gene expression data. Bioinformatics 26, 139–140. 10.1093/bioinformatics/btp616.

114. Kolmogorov, M., Armstrong, J., Raney, B.J., Streeter, I., Dunn, M., Yang, F., Odom, D., Flicek, P., Keane, T.M., Thybert, D., et al. (2018). Chromosome assembly of large and complex genomes using multiple references. Genome Res. 28, 1720–1732. 10.1101/gr.236273.118.

115. Kolmogorov, M., Raney, B., Paten, B., and Pham, S. (2014). Ragout—a reference-assisted assembly tool for bacterial genomes. Bioinformatics 30, i302–i309. 10.1093/bioinformatics/btu280.

116. Katoh, K., and Standley, D.M. (2013). MAFFT multiple sequence alignment software version 7: improvements in performance and usability. Mol Biol Evol 30, 772–780. 10.1093/molbev/mst010.

117. Robinson, J.T., Thorvaldsdóttir, H., Winckler, W., Guttman, M., Lander, E.S., Getz, G., and Mesirov, J.P. (2011). Integrative genomics viewer. Nat Biotechnol 29, 24–26. 10.1038/nbt.1754.

118. Stamatakis, A. (2014). RAxML version 8: a tool for phylogenetic analysis and post-analysis of large phylogenies. Bioinformatics 30, 1312–1313. 10.1093/bioinformatics/btu033.

119. Suchard, M.A., Lemey, P., Baele, G., Ayres, D.L., Drummond, A.J., and Rambaut, A. (2018). Bayesian phylogenetic and phylodynamic data integration using BEAST 1.10. Virus Evolution 4. 10.1093/ve/vey016.

120. Waterhouse, A.M., Procter, J.B., Martin, D.M.A., Clamp, M., and Barton, G.J. (2009). Jalview Version 2--a multiple sequence alignment editor and analysis workbench. Bioinformatics 25, 1189–1191. 10.1093/bioinformatics/btp033.

121. Jumper, J., Evans, R., Pritzel, A., Green, T., Figurnov, M., Ronneberger, O., Tunyasuvunakool, K., Bates, R., Žídek, A., Potapenko, A., et al. (2021). Highly accurate protein structure prediction with AlphaFold. Nature 596, 583–589. 10.1038/s41586-021-03819-2.

122. Meng, E.C., Goddard, T.D., Pettersen, E.F., Couch, G.S., Pearson, Z.J., Morris, J.H., and Ferrin, T.E. (2023). UCSF C_HIMERA_X1: Tools for structure building and analysis. Protein Science 32, e4792. 10.1002/pro.4792.

123. Wick, R.R., Judd, L.M., Gorrie, C.L., and Holt, K.E. (2017). Unicycler: Resolving bacterial genome assemblies from short and long sequencing reads. PLoS Comput Biol 13, e1005595. 10.1371/journal.pcbi.1005595.

124. Wick, R.R., Schultz, M.B., Zobel, J., and Holt, K.E. (2015). Bandage: interactive visualization of *de novo* genome assemblies. Bioinformatics 31, 3350–3352. 10.1093/bioinformatics/btv383.

125. Hyatt, D., Chen, G.-L., LoCascio, P.F., Land, M.L., Larimer, F.W., and Hauser, L.J. (2010). Prodigal: prokaryotic gene recognition and translation initiation site identification. BMC Bioinformatics 11, 119. 10.1186/1471-2105-11-119.

126. Seemann, T. (2014). Prokka: rapid prokaryotic genome annotation. Bioinformatics 30, 2068–2069. 10.1093/bioinformatics/btu153.

127. Feng, Q., Liang, S., Jia, H., Stadlmayr, A., Tang, L., Lan, Z., Zhang, D., Xia, H., Xu, X., Jie, Z., et al. (2015). Gut microbiome development along the colorectal adenoma-carcinoma sequence. Nat Commun 6, 6528. 10.1038/ncomms7528.

128. David, L.A., Weil, A., Ryan, E.T., Calderwood, S.B., Harris, J.B., Chowdhury, F., Begum, Y., Qadri, F., LaRocque, R.C., and Turnbaugh, P.J. (2015). Gut microbial succession follows acute secretory diarrhea in humans. mBio 6, e00381–00315. 10.1128/mBio.00381-15.

129. De Carvalho, F.M., Valiatti, T.B., Santos, F.F., Silveira, A.C.D.O., Guimarães, A.P.C., Gerber, A.L., Souza, C.D.O., Cassu Corsi, D., Brasiliense, D.M., Castelo-Branco, D.D.S.C.M., et al. (2022). Exploring the Bacteriome and Resistome of Humans and Food-Producing Animals in Brazil. Microbiol Spectr 10, e00565–22. 10.1128/spectrum.00565-22.

130. Raymond, F., Ouameur, A.A., Déraspe, M., Iqbal, N., Gingras, H., Dridi, B., Leprohon, P., Plante, P.-L., Giroux, R., Bérubé, È., et al. (2016). The initial state of the human gut microbiome determines its reshaping by antibiotics. ISME J 10, 707–720. 10.1038/ismej.2015.148.

131. Qin, J., Li, Y., Cai, Z., Li, S., Zhu, J., Zhang, F., Liang, S., Zhang, W., Guan, Y., Shen, D., et al. (2012). A metagenome-wide association study of gut microbiota in type 2 diabetes. Nature 490, 55–60. 10.1038/nature11450.

132. Wen, C., Zheng, Z., Shao, T., Liu, L., Xie, Z., Le Chatelier, E., He, Z., Zhong, W., Fan, Y., Zhang, L., et al. (2017). Quantitative metagenomics reveals unique gut microbiome biomarkers in ankylosing spondylitis. Genome Biol 18, 142. 10.1186/s13059-017-1271-6.

133. Le Chatelier, E., Nielsen, T., Qin, J., Prifti, E., Hildebrand, F., Falony, G., Almeida, M., Arumugam, M., Batto, J.-M., Kennedy, S., et al. (2013). Richness of human gut microbiome correlates with metabolic markers. Nature 500, 541–546. 10.1038/nature12506.

134. Xie, H., Guo, R., Zhong, H., Feng, Q., Lan, Z., Qin, B., Ward, K.J., Jackson, M.A., Xia, Y., Chen, X., et al. (2016). Shotgun Metagenomics of 250 Adult Twins Reveals Genetic and Environmental Impacts on the Gut Microbiome. Cell Syst 3, 572–584.e3. 10.1016/j.cels.2016.10.004.

135. Yassour, M., Jason, E., Hogstrom, L.J., Arthur, T.D., Tripathi, S., Siljander, H., Selvenius, J., Oikarinen, S., Hyöty, H., Virtanen, S.M., et al. (2018). Strain-Level Analysis of Mother-to-Child Bacterial Transmission during the First Few Months of Life. Cell Host Microbe 24, 146–154.e4. 10.1016/j.chom.2018.06.007.

136. Dhakan, D.B., Maji, A., Sharma, A.K., Saxena, R., Pulikkan, J., Grace, T., Gomez, A., Scaria, J., Amato, K.R., and Sharma, V.K. (2019). The unique composition of Indian gut microbiome, gene catalogue, and associated fecal metabolome deciphered using multi-omics approaches. Gigascience 8, giz004. 10.1093/gigascience/giz004.

137. Zeevi, D., Korem, T., Zmora, N., Israeli, D., Rothschild, D., Weinberger, A., Ben-Yacov, O., Lador, D., Avnit-Sagi, T., Lotan-Pompan, M., et al. (2015). Personalized Nutrition by Prediction of Glycemic Responses. Cell 163, 1079–1094. 10.1016/j.cell.2015.11.001.

138. Ferretti, P., Pasolli, E., Tett, A., Asnicar, F., Gorfer, V., Fedi, S., Armanini, F., Truong, D.T., Manara, S., Zolfo, M., et al. (2018). Mother-to-Infant Microbial Transmission from Different Body Sites Shapes the Developing Infant Gut Microbiome. Cell Host Microbe 24, 133–145.e5. 10.1016/j.chom.2018.06.005.

139. Kim, C.Y., Lee, M., Yang, S., Kim, K., Yong, D., Kim, H.R., and Lee, I. (2021). Human reference gut microbiome catalog including newly assembled genomes from under-represented Asian metagenomes. Genome Med 13, 134. 10.1186/s13073-021-00950-7.

140. Liu, W., Zhang, J., Wu, C., Cai, S., Huang, W., Chen, J., Xi, X., Liang, Z., Hou, Q., Zhou, B., et al. (2016). Unique Features of Ethnic Mongolian Gut Microbiome revealed by metagenomic analysis. Sci Rep 6, 34826. 10.1038/srep34826.

141. Zhernakova, A., Kurilshikov, A., Bonder, M.J., Tigchelaar, E.F., Schirmer, M., Vatanen, T., Mujagic, Z., Vila, A.V., Falony, G., Vieira-Silva, S., et al. (2016). Population-based metagenomics analysis reveals markers for gut microbiome composition and diversity. Science 352, 565–569. 10.1126/science.aad3369.

142. Obregon-Tito, A.J., Tito, R.Y., Metcalf, J., Sankaranarayanan, K., Clemente, J.C., Ursell, L.K., Zech Xu, Z., Van Treuren, W., Knight, R., Gaffney, P.M., et al. (2015). Subsistence strategies in traditional societies distinguish gut microbiomes. Nat Commun 6, 6505. 10.1038/ncomms7505.

143. Li, J., Jia, H., Cai, X., Zhong, H., Feng, Q., Sunagawa, S., Arumugam, M., Kultima, J.R., Prifti, E., Nielsen, T., et al. (2014). An integrated catalog of reference genes in the human gut microbiome. Nat Biotechnol 32, 834–841. 10.1038/nbt.2942.

144. Bäckhed, F., Roswall, J., Peng, Y., Feng, Q., Jia, H., Kovatcheva-Datchary, P., Li, Y., Xia, Y., Xie, H., Zhong, H., et al. (2015). Dynamics and Stabilization of the Human Gut Microbiome during the First Year of Life. Cell Host Microbe 17, 690–703. 10.1016/j.chom.2015.04.004.

145. Karlsson, F.H., Tremaroli, V., Nookaew, I., Bergström, G., Behre, C.J., Fagerberg, B., Nielsen, J., and Bäckhed, F. (2013). Gut metagenome in European women with normal, impaired and diabetic glucose control. Nature 498, 99–103. 10.1038/nature12198.

146. Lou, Y.C., Olm, M.R., Diamond, S., Crits-Christoph, A., Firek, B.A., Baker, R., Morowitz, M.J., and Banfield, J.F. (2021). Infant gut strain persistence is associated with maternal origin, phylogeny, and traits including surface adhesion and iron acquisition. Cell Rep Med 2, 100393. 10.1016/j.xcrm.2021.100393.

147. Human Microbiome Project Consortium (2012). A framework for human microbiome research. Nature 486, 215–221. 10.1038/nature11209.

148. Bittinger, K., Zhao, C., Li, Y., Ford, E., Friedman, E.S., Ni, J., Kulkarni, C.V., Cai, J., Tian, Y., Liu, Q., et al. (2020). Bacterial colonization reprograms the neonatal gut metabolome. Nat Microbiol 5, 838– 847. 10.1038/s41564-020-0694-0.

149. Eren, A.M., Kiefl, E., Shaiber, A., Veseli, I., Miller, S.E., Schechter, M.S., Fink, I., Pan, J.N., Yousef, M., Fogarty, E.C., et al. (2020). Community-led, integrated, reproducible multi-omics with anvi’o. Nat Microbiol 6, 3–6. 10.1038/s41564-020-00834-3.

150. Eren, A.M., Vineis, J.H., Morrison, H.G., and Sogin, M.L. (2013). A Filtering Method to Generate High Quality Short Reads Using Illumina Paired-End Technology. PLoS ONE 8, e66643. 10.1371/journal.pone.0066643.

151. Danecek, P., Bonfield, J.K., Liddle, J., Marshall, J., Ohan, V., Pollard, M.O., Whitwham, A., Keane, T., McCarthy, S.A., Davies, R.M., et al. (2021). Twelve years of SAMtools and BCFtools. Gigascience 10, giab008. 10.1093/gigascience/giab008.

152. Wood, D.E., Lu, J., and Langmead, B. (2019). Improved metagenomic analysis with Kraken 2. Genome Biol 20, 257. 10.1186/s13059-019-1891-0.

