## Supplementary figures and images for "A pervasive large conjugative plasmid mediates multispecies biofilm formation in the intestinal microbiota increasing resilience to perturbations"

### Supplemental Figure 1

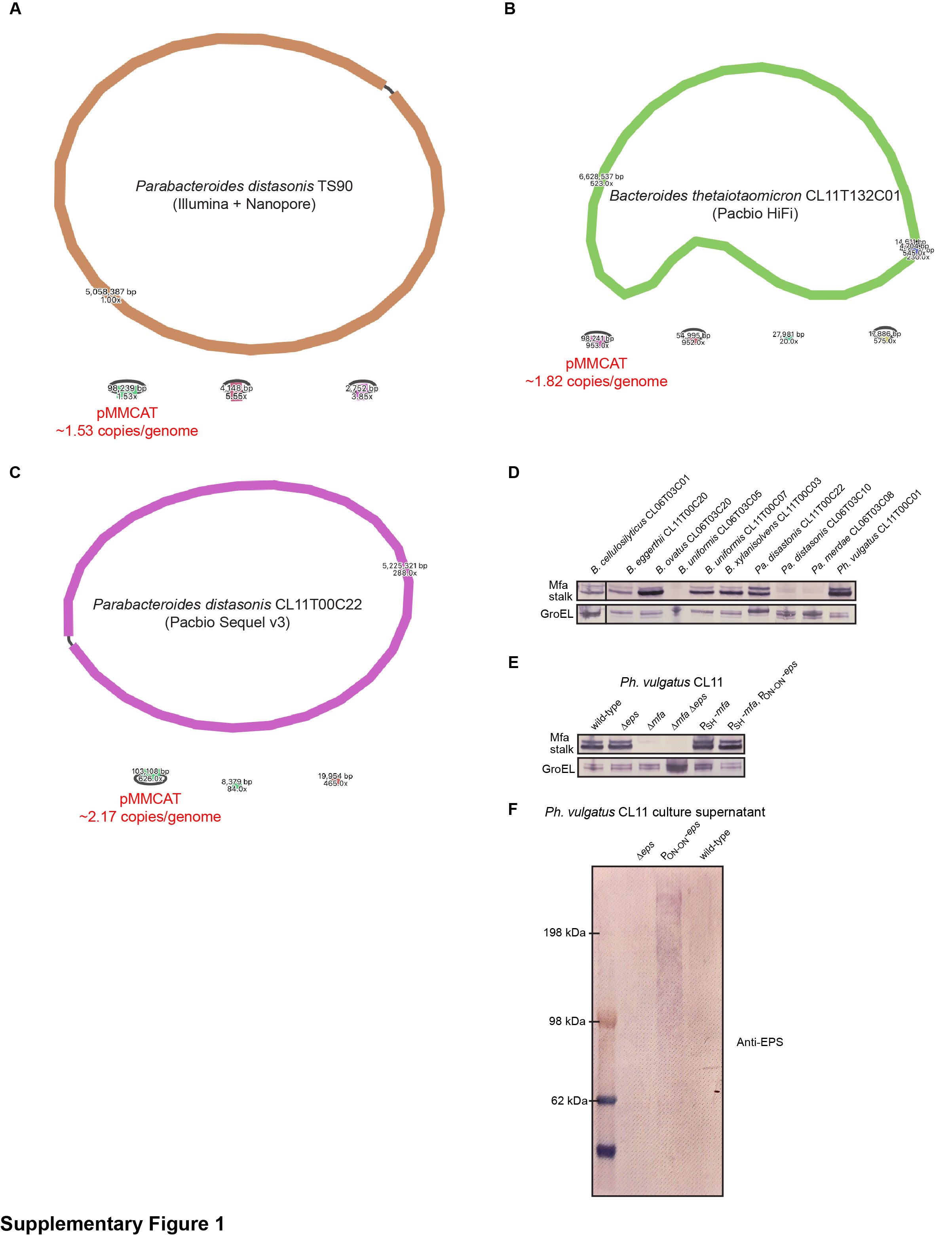

### Supplemental Figure 3

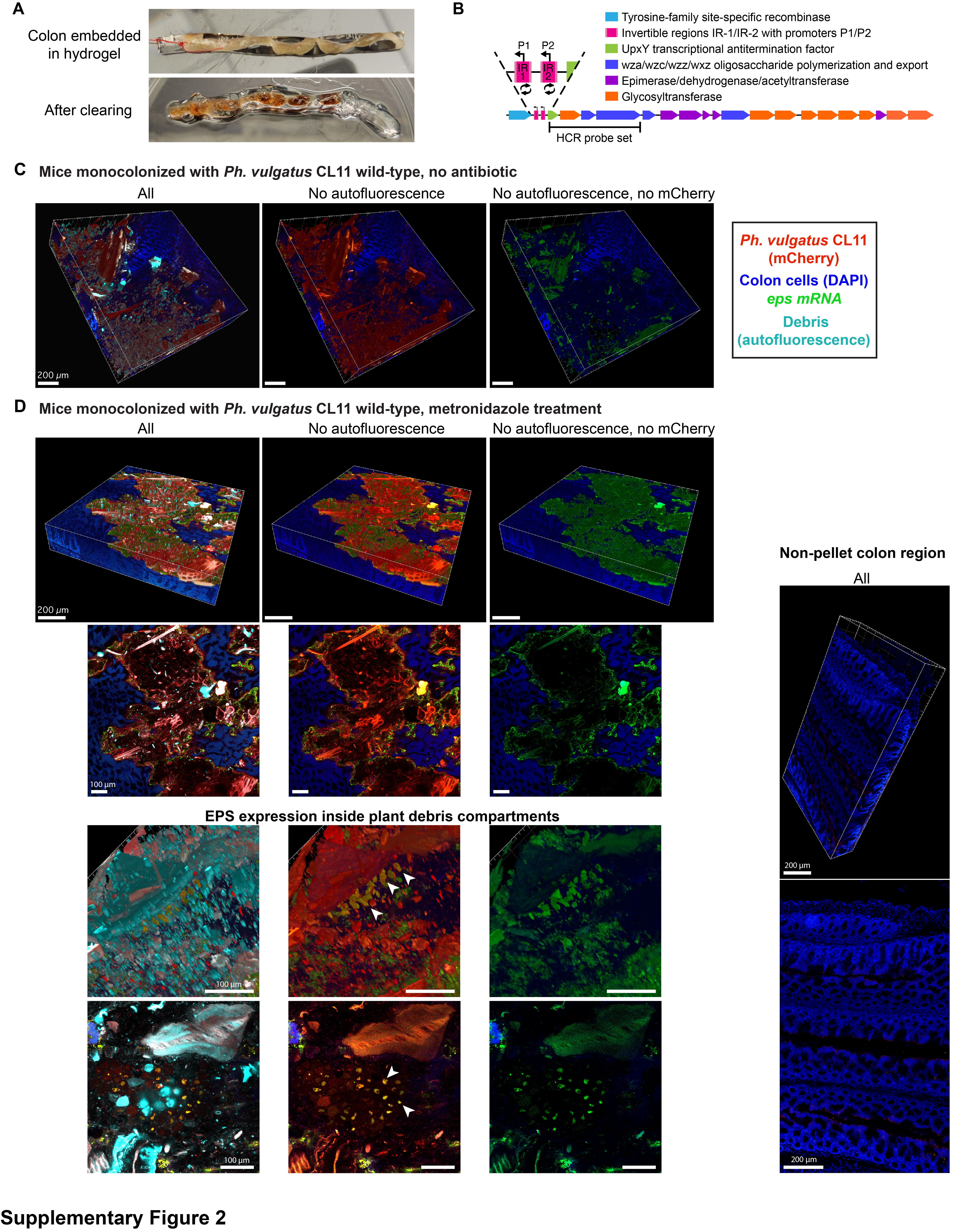

### Supplemental Figure 3

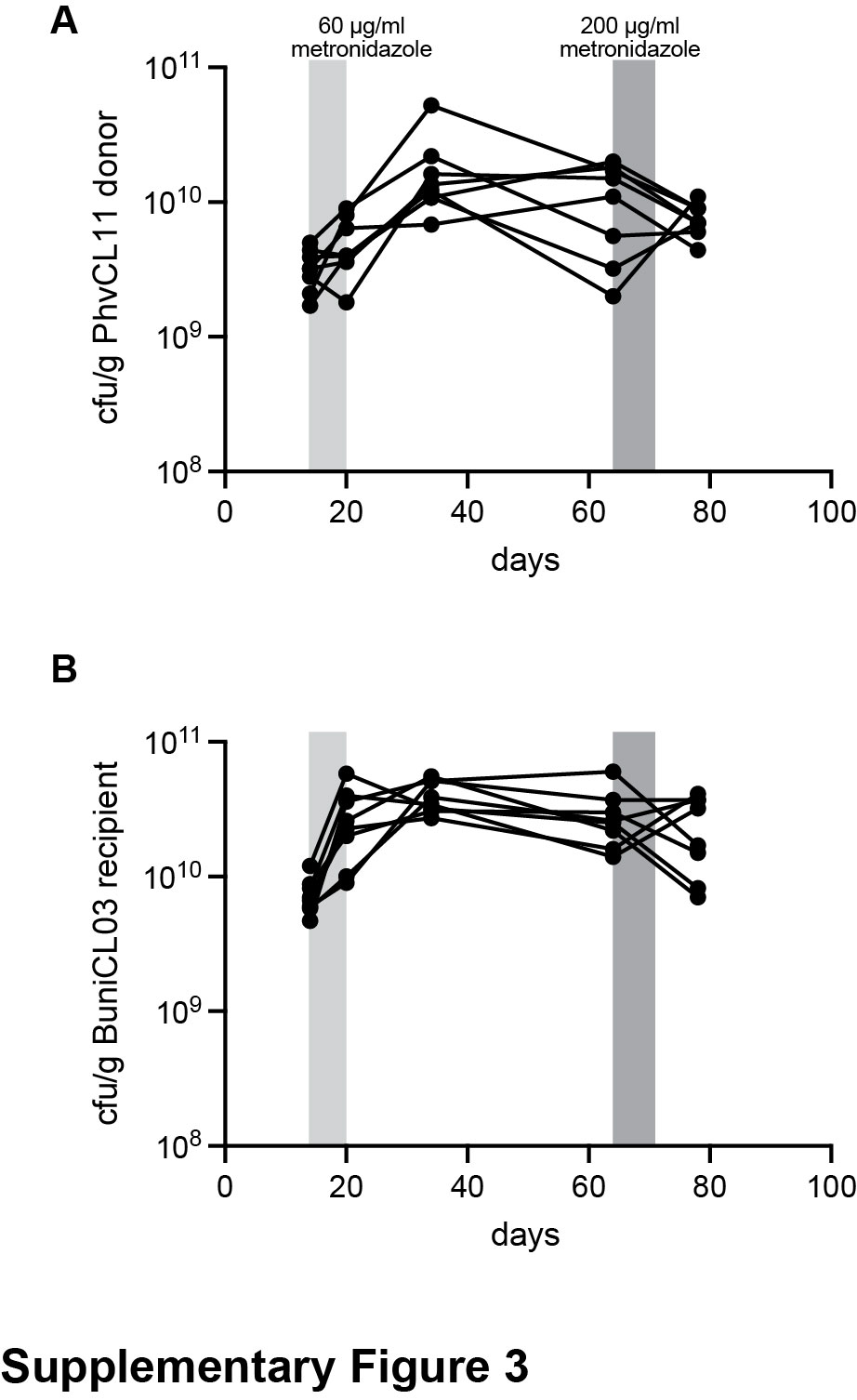

### Supplemental Figure 4

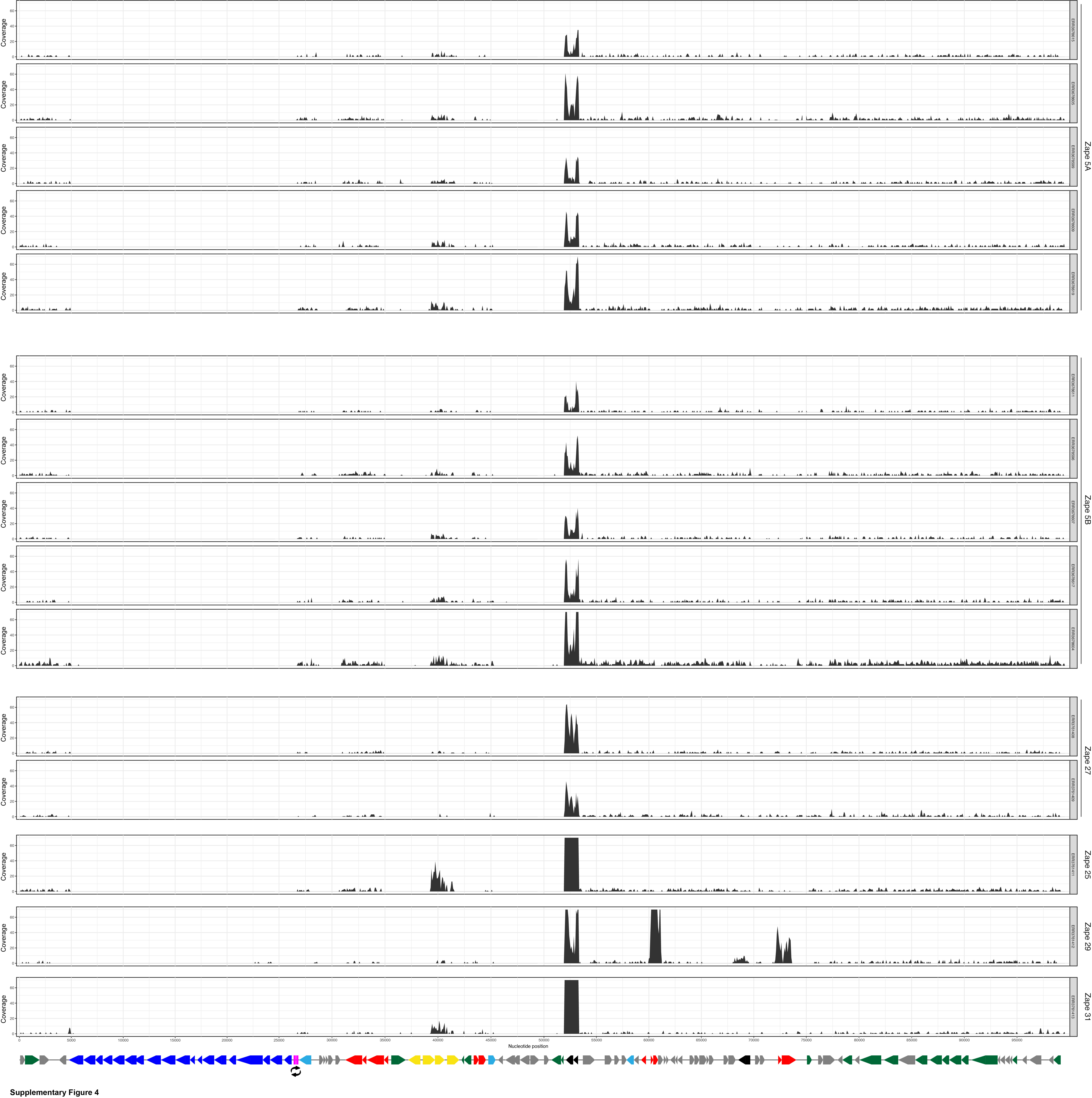

### Supplemental Figure 5

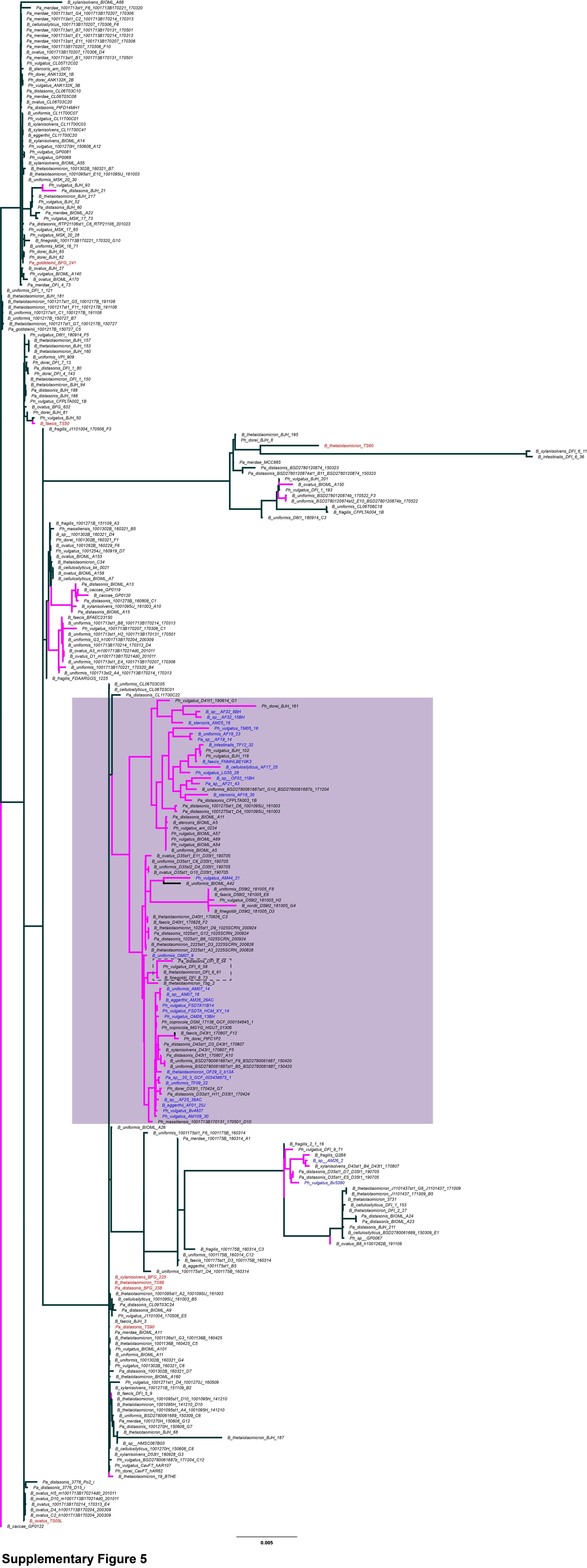

### Supplemental Figure 6

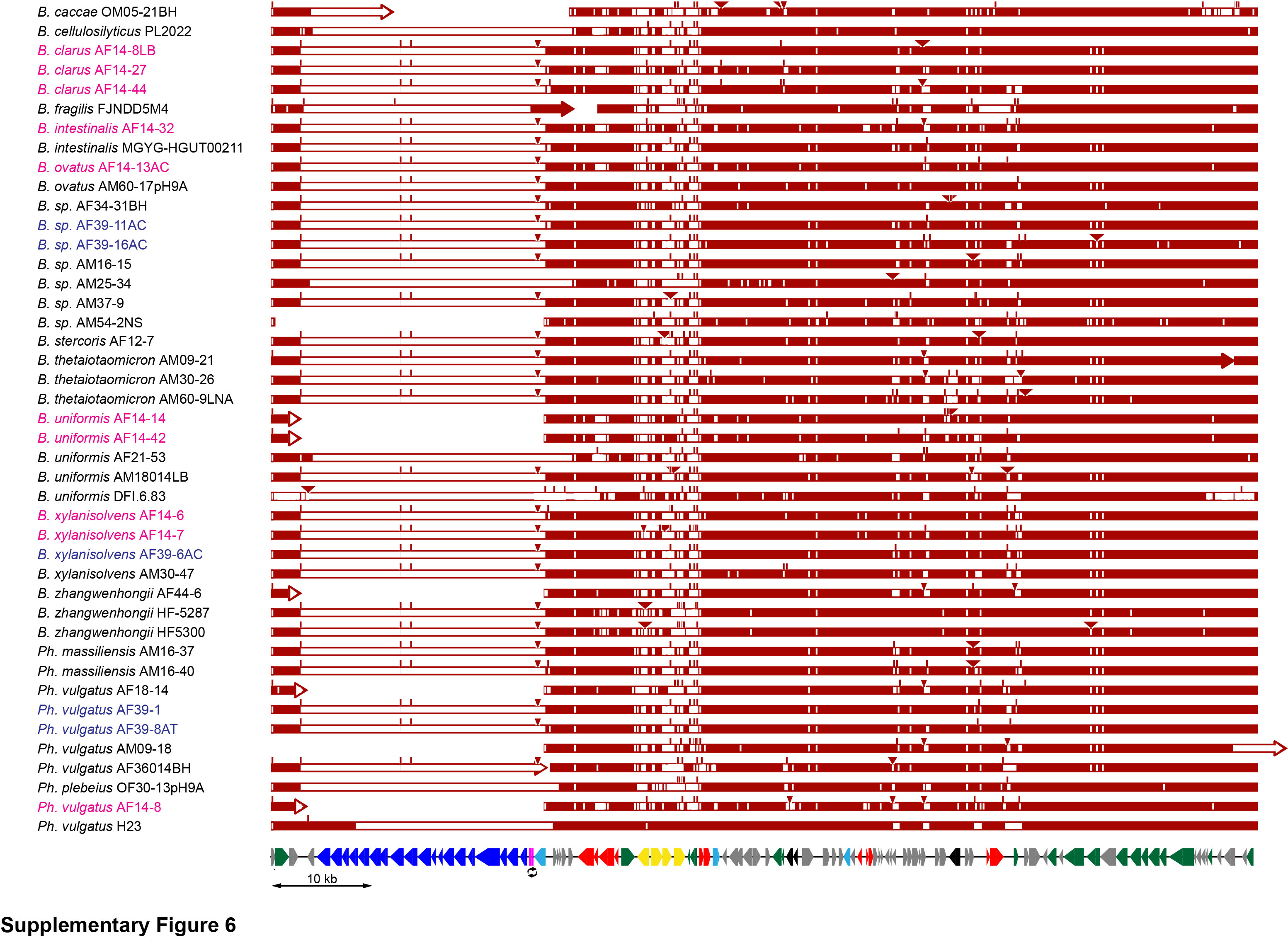

### Supplemental Figure 7

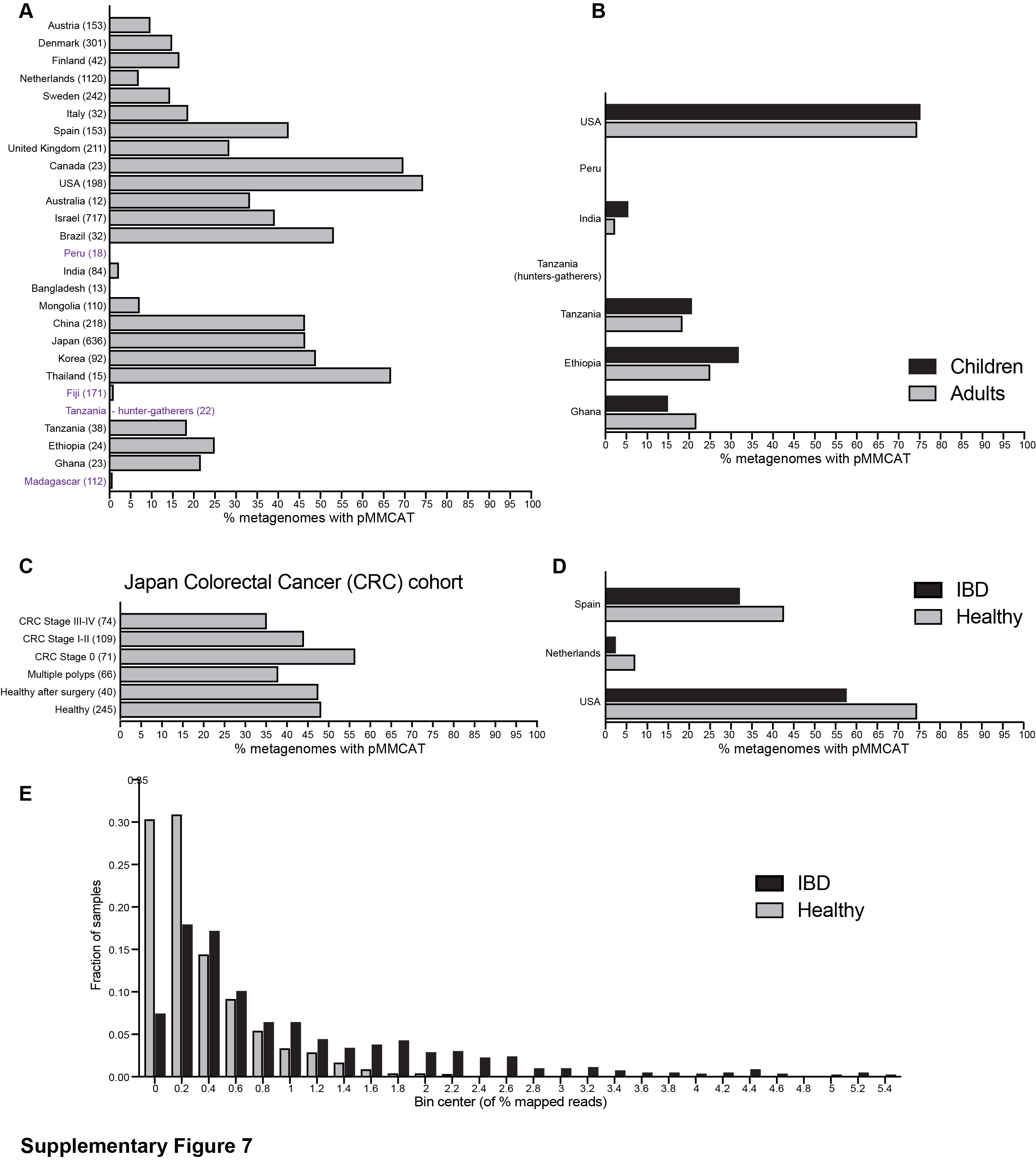
